# Drosophila species learn dialects through communal living

**DOI:** 10.1101/206920

**Authors:** Balint Z Kacsoh, Julianna Bozler, Giovanni Bosco

## Abstract

Many species are able to share information about their environment by communicating through auditory, visual, and olfactory cues. In Drosophila melanogaster, exposure to parasitoid wasps leads to a decline in egg laying, and exposed females communicate this threat to naïve flies, which also depress egg laying. We find that species across the genus Drosophila respond to wasps by egg laying reduction, activate cleaved caspase in oocytes, and communicate the presence of wasps to naïve individuals. communication within a species and between closely related species is efficient, while more distantly related species exhibit partial communication. Remarkably, partial communication between some species is enhanced after a cohabitation period that requires exchange of visual and olfactory signals. This interspecies “dialect learning” requires neuronal cAMP signaling in the mushroom body, suggesting neuronal plasticity facilitates dialect learning and memory. These observations establish Drosophila as genetic models for inter-species social communication and evolution of dialects.

---

The ability to interpret environmental information is a phenomenon found throughout all life forms. From bacteria to plants and to mammals, communication occurs within as well as between species. In some cases, information that is being shared can be highly specific, such as in the case of honeybees communicating instructions on where to find nectar (1–3). In other cases, opportunistic bystanders can also benefit from general information. For example, predator alarm calls generated as a warning are observed, where multiple species participate in repeating the alarm throughout the community (4–8). In all cases, the information that is shared can be dependent on local environmental cues and experiences and the manner in which information is communicated is strongly influenced by past experiences of each individual. For example, birds, which live in geographically distinct populations, manifest unique song variants or regional dialects that can last for decades, but these animals are nevertheless still able to communicate with others of their species (9–11). Because dialects are learned and therefore influenced (12) by specific local environmental differences, it suggests that both social and non-social experiences can have dramatic effects on cognitive development (13).

It is proposed that a myriad of environmental cues, both social and non-social, are critical to animal development in determining the ability to convey and receive specific types of information. However, there are many outlying questions as a result of this proposition: What cues are important? When are these cues important? How can environmental cues interact with genetically determined developmental programs? Although social communication is most extensively documented in more derived organisms such as mammals and birds, insects can also display a broad range of behavioral tasks. Bees are known to be able to learn from non-natural sources in order to obtain a reward through social learning. Such information can be passed on to naïve, student bees through the use of visual cues (14,15). Insect social learning extends to the genetic model system of Drosophila, where student, observer flies learn from a trained, teacher-fly, using visual cues. This has been shown in communication involving food sources and predator threats (16,17).

Chemical cues can serve as intra- and inter-species signals, such as fox and guinea pig urine affecting not only conspecific behavior, but also the behavior of other animals (18–20). Sound can also be used, such as in bats and bottlenose dolphins, which are able to distinguish members of the community through the use of echolocation pitch recognition (21,22). Plants have a vast arsenal of responses to pathogens (23), including communicating a threat to neighboring plants through the use of volatile organic compounds (24). Plant inter-species (25-31) and intra-species (32–34) communication occurs both in laboratory settings and in the wild (30,35).

*Drosophila melanogaster* and other Drosophila species have provided insights into mechanisms of learning, memory, and complex behaviors (36,37). However, these behaviors and phenotypes have been studied almost exclusively in domesticated *D. melanogaster* lab monocultures, while *D. melanogaster* wild populations are surrounded by a broad range of predators, microbes, and other Drosophilids, highlighting a communal component of the organism’s life cycle (38). This raises the possibility of behavioral phenomenon that have yet to be discovered and analyzed in domesticated lab monocultures (39–41). Given the vast range of environmental inputs on a wild Drosophilid, a fly must be able to discern important information from extraneous inputs, while interacting with conspecifics and a variety of other species.

Although modes of intra- and inter-species communication are likely to be genetically limited, there is also value in learning to interpret signals from variable, local environments that may provide immediate survival benefits. How do genetically constrained neurological features and variable environmental factors interact to produce context-dependent, meaningful information? Under which environmental factors would information sharing between different species occur and be beneficial? In this study, we sought to begin to address these questions in the Drosophila model system by using a pan-Drosophila predator known to elicit social communication (17,42). *D. melanogaster* presented with parasitoid wasps have multiple behavioral responses, including a reduction in oviposition (egg laying) through an increase in ovarian apoptosis (17,43–46). After removal of the wasp, a wasp-exposed “teacher” fly can instruct a naïve “student” fly about the presence of the wasp threat through the exclusive use of visual cues, such that students now reduce their own oviposition by triggering ovarian apoptosis. Using this fly-fly social communication paradigm we asked (1) whether social communication is conserved among other Drosophila species, (2) if Drosophilids engage in interspecies communication, and (3) what environmental and genetic factors are required for interspecies communication.

## RESULTS

### INTRA- AND INTER-SPECIES COMMUNICATION

We utilized the fly duplex, an apparatus with two transparent acrylic compartments to test whether different species respond to seeing predators (acute response) and if exposed “teacher” female flies can communicate this threat to naïve unexposed “student” female flies (17). The duplex allows flies to see other flies or wasps in the adjacent compartment, without direct contact, making all communication only visual (Fig. 1A). Ten female and two male flies are placed into one duplex compartment, with an adjacent compartment containing twenty female wasps. Following a 24-hour exposure, wasps are removed and acute response is measured by counting the number of eggs laid in the first 24-hour period in a blinded manner. Flies are shifted to a new duplex, with ten female and two male naïve student flies in the adjacent compartment (Fig. 1 A, see methods). Following a second 24-hour period, all flies are removed and the response of both teacher and student is measured by counting the number of eggs laid in a blinded manner. The 24-48-hour period measures memory of teachers having seen the wasps and students having learned from the teachers. Using wild-type *D. melanogaster*, we find both an acute response and a memory response to the wasp in teacher flies and a learned response in naïve student flies (Fig. 1 B, Fig. S 1 A) (17,45,46).

**Figure 1.**
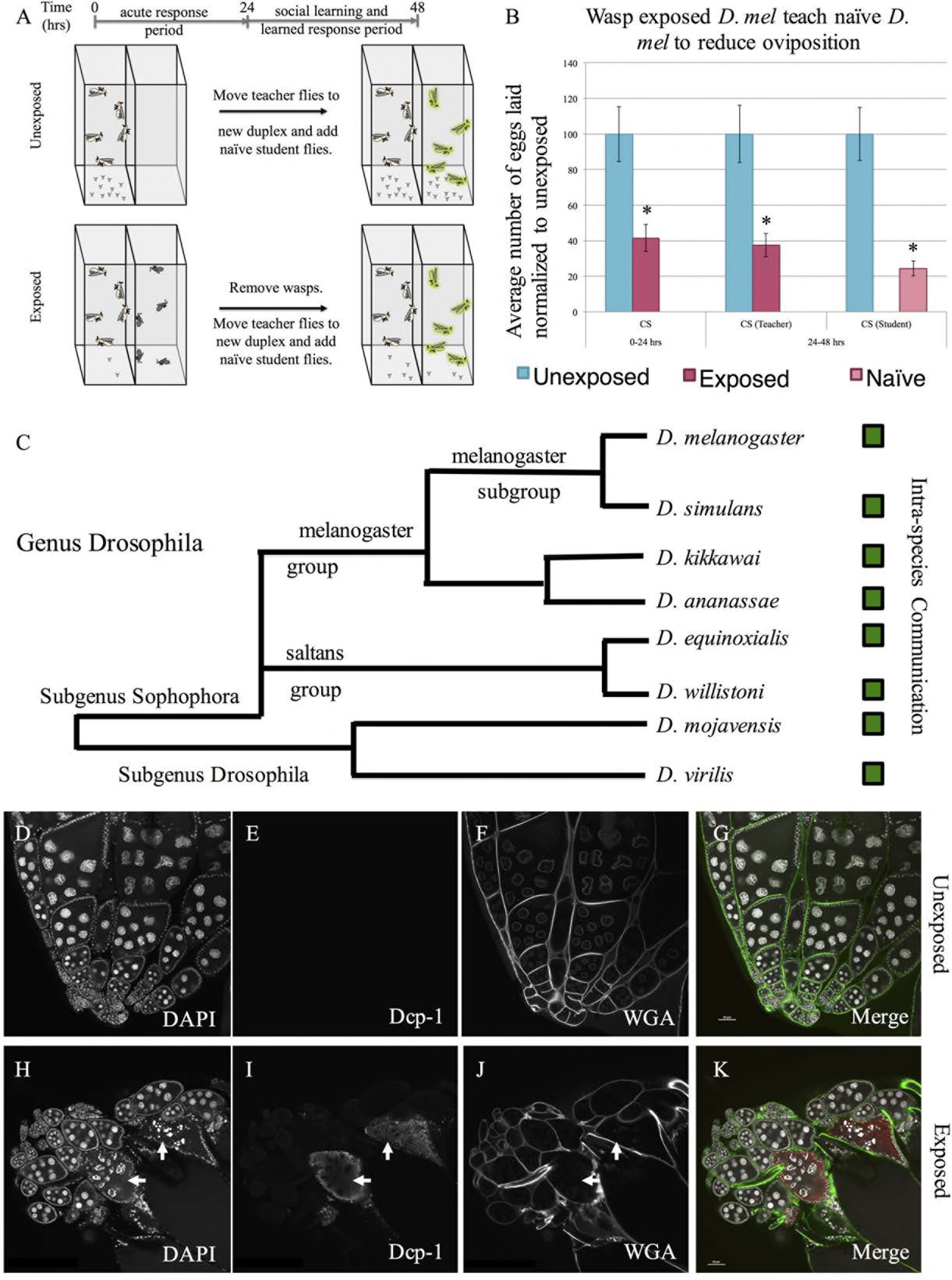
A predator threat is communicated through visual cues within species across the genus Drosophila, modulating reproductive behavior and caspase activation. (A) Standard experimental design. (B) Percentage of eggs laid by exposed flies normalized to eggs laid by unexposed flies is shown. Wild-type *D. melanogaster* (Canton S) exposed to wasps lay fewer eggs than unexposed flies. (C) Phylogeny of 8 species tested across the genus Drosophila that demonstrate the ability to communicate through visual cues. Green boxes indicate social learning is present in species tested. Representative ovary of control and wasp exposed Drosophila showing caspase activation (*D. melanogaster*). DAPI (D, H), activated Dcp-1 (E, I), WGA (F,J), and the merged images (G, K) are shown. Arrows denote apoptotic egg chambers. Error bars represent standard error (n = 12 biological replicates) (*p < 0.05).

We then asked whether the acute, memory, and student social learning behaviors are conserved in other Drosophila species, with varying relatedness to *D. melanogaster* ranging from sister species, such as *D. simulans*, to very distantly related species, such as *D. virilis*. For each species, we tested a sister species as an additional way to validate our observations. Across a broad span of the genus Drosophila, we find the conservation of both the acute and memory responses in teacher flies in addition to the ability of teachers to communicate to student flies. (Fig. 1 C, Fig. S 1 B-H). Some of these species have been previously shown to depress oviposition during wasp exposure (46). Our experimental design allows for only visual cues to be detected from the wasps and from teachers to student flies. Thus, in all species tested, visual cues are sufficient for flies to detect wasps and for naïve flies to learn from wasp-exposed teacher flies. Conservation of these behaviors is especially impressive as the species tested are separated by millions of years of evolution, yet the basic behaviors observed in *D. melanogaster* are maintained. Moreover, this conservation further underscores the importance this innate behavior must have since even laboratory cultures that have not experienced wasp for many generations nevertheless exhibit a robust response.

Oviposition reduction is modulated in part by the effector caspase Dcp-1 (17). In *D. melanogaster*, we observe overlapping staining of activated Dcp-1 with a punctate pattern of DNA staining with 4’, 6-diamidino-2-phenylindole (DAPI), indicative of oocyte specific apoptotic activity (Fig. 1 D-K, Fig. S 2). We performed immunofluorescence of activated Dcp-1 across a broad range of Drosophila species, revealing cleaved caspase following wasp exposure in all 15 Drosophila species tested (Fig. S 3). We demonstrate an increase in positive cleaved caspase oocytes following wasp exposure (Fig. S 4), along with a decrease in total number of egg chambers (Fig. S 5), suggestive of ovarian apoptosis and elimination of oocytes (17). Phylogenetic trees shown are adapted from previous work (47).

Following the observation that an acute response to wasps and intra-species communication is conserved across the genus, we asked whether the wasp threat could be communicated between two different species. We utilized 15 Drosophila species that respond to wasps to answer this question (Fig. S 3,4). The species were selected to span the phylogeny with different degrees of relatedness to *D. melanogaster* (47). We find that *D. melanogaster* are able to communicate the threat to and receive communications from closely related species, such as *D. simulans* and *D. yakuba*, with oviposition of students paired with wasp-exposed teachers being ~10-30% compared to unexposed (Fig. 2 A-B, Fig. S 6 A-F). Interestingly, species more distantly related to *D. melanogaster*, such as *D. ananassae* and its sister species, elicit a partial communication phenotype, with oviposition depression of students paired with wasp-exposed teachers being ~50-65% of unexposed flies (Fig. 2 C-F, Fig. S 6 G-J). A second strain isolate of *D. ananassae* also show partial communication with *D. melanogaster* (Fig. 2 C-F, Fig. S 6 G-J). Species more distantly related to *D. melanogaster*, such as *D. willistoni* and *D. virilis*, cannot communicate with *D. melanogaster* (Fig. 2 G-J, Fig. S 6 K-P). Collectively, the data suggest that evolutionary distance contributes the to the efficiency of interspecies communication. *D. ananassae* show varying communication phenotypes with other Drosophila species, though the pattern of communication is different. For example, *D. ananassae* exhibit partial communication with *D. simulans* (Fig. S 7 A-B), strong communication with its sister *D. kikkawai* (Fig. S 7 CD), and partial communication with *D. equinoxialis* and *D. willistoni* (Fig. S 7 E-H). D. *ananassae*, in addition to *D. melanogaster*, are unable to communicate with the distantly related *D. mojavensis* and *D. virilis* (Fig. 2 I-J, Fig. S 7 I-L). Species such as *D. virilis*, which were unable to communicate with *D. melanogaster* and *D. ananassae*, can communicate with other species, such as its sister species *D. mojavensis* (Fig. 2 K-L). Thus, although all species tested are capable of intra-species communication, there is a fundamental, species-specific difference in “fly language” or in strategy for communicating wasp threat.

**Figure 2.**
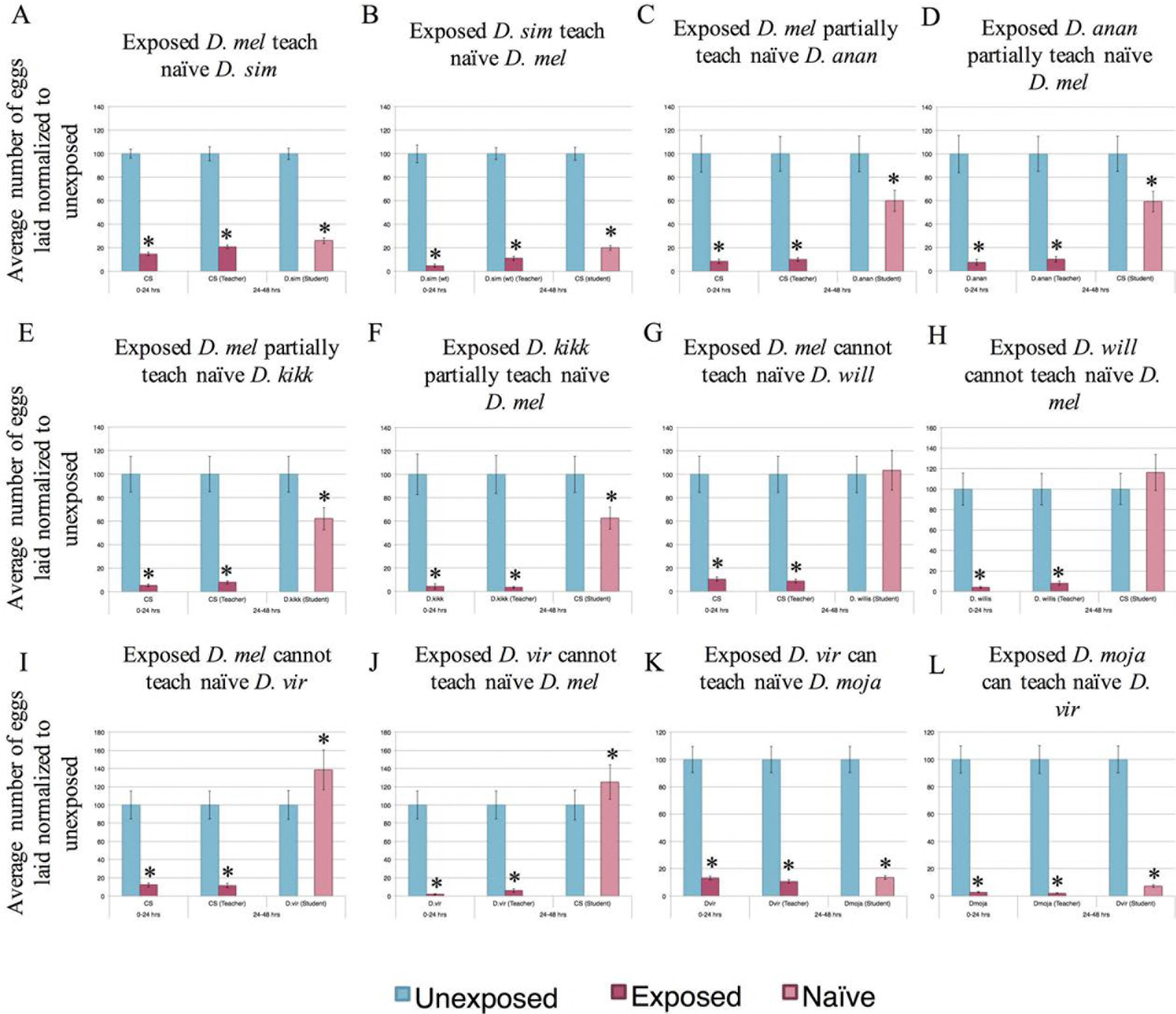
Interspecies communication of predator threats. Percentage of eggs laid by exposed flies normalized to eggs laid by unexposed flies is shown. Flies exposed to wasps lay fewer eggs than unexposed flies. Communication between *D. melanogaster* and: *D. simulans* (A, B), *D. ananassae* (C, D), *D. kikkawai* (E,F), *D. willistoni* (G,H), and *D. virilis* (I,J) shows varying communication abilities.. Communication between *D. virilis* and *D. mojavensis* occurs (K,L). Error bars represent standard error (n = 12 biological replicates) (*p < 0.05).

### DIALECT LEARNING

Given that closely related species can communicate the threat of a wasp, we examined the environmental factors that could be contributing to such interspecies communication. Specifically, we tested whether a period of cohabitation with frequent contact between two poorly communicating species could improve interspecies communication. *D. melanogaster* were cohabitated with species capable of only partial communication (*e.g. D. ananassae*) (Fig. 2 C-D) for one week in a single container, allowing for frequent and multiple channels of sensory interactions. Following a weeklong cohabitation period, the two species were separated and used as students paired with teachers of the other species (Fig. 3 A). In all experiments teachers had existed only as a monoculture, while all flies experiencing an interspecies cohabitation period were subsequently used only as students.

**Figure 3.**
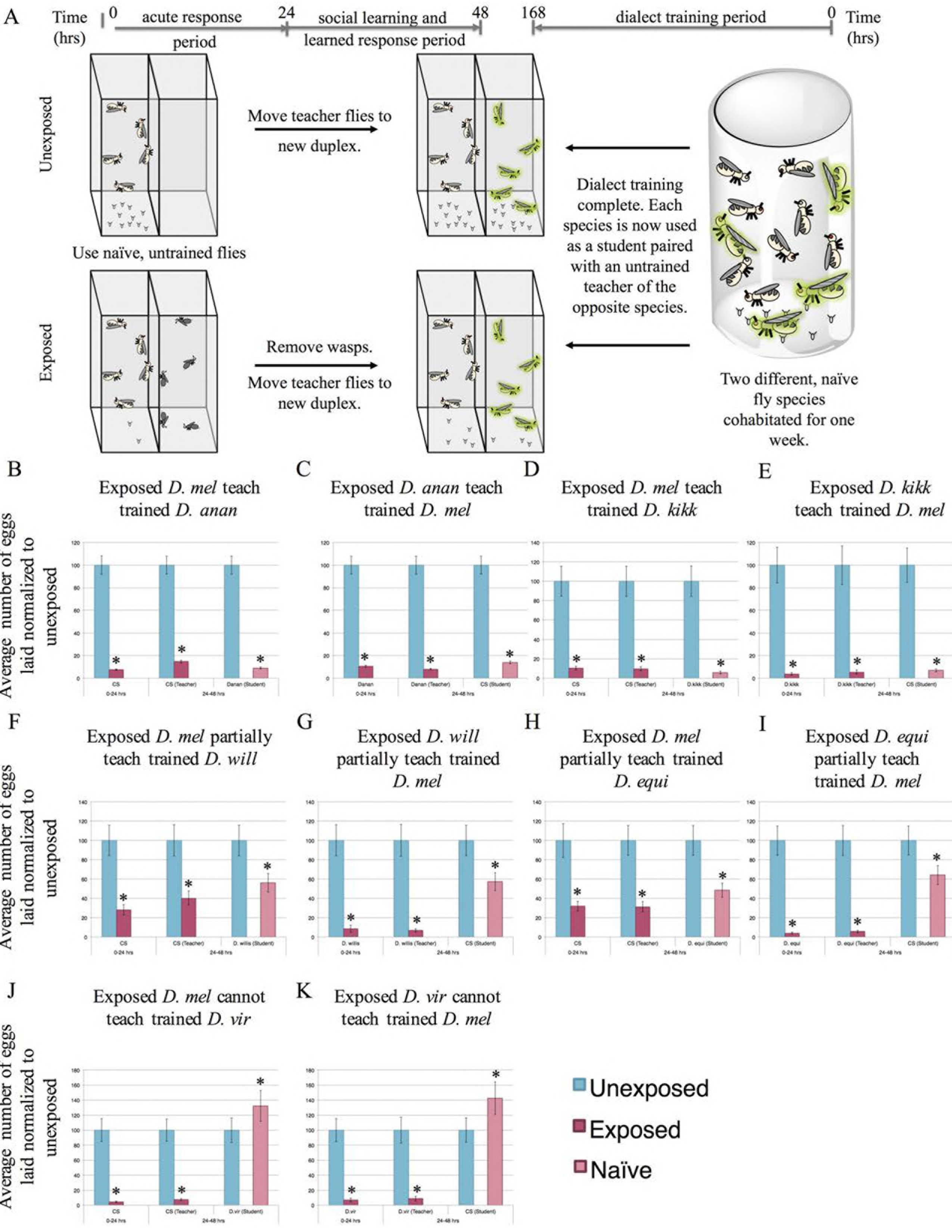
Species cohabitation enables inter-species communication. (A) Experimental design of dialect training for flies that are used as students. Two species are cohabitated for one week prior to being used as students for naive, untrained teacher flies of the opposite species. Percentage of eggs laid by exposed flies normalized to eggs laid by unexposed flies is shown. Communication between trained students *D. melanogaster* and: *D. ananassae* showing strong communication following cohabitation (B, C), *D. kikkawai* showing strong communication following co-incubation (D, E), *D. willistoni* showing partial communication following co-incubation (F,G), *D. equinoxialis* showing partial communication following coincubation (H,I), and *D. virilis* showing no communication following co-incubation (J,K). Error bars represent standard error (n = 12 biological replicates) (*p < 0.05).

We find that cohabitation can greatly enhance communication between some species, suggesting that some form of training occurs during this period. After cohabitation, *D. ananassae* learn very efficiently from *D. melanogaster* teachers, demonstrating that cohabitation of two species yields an expanded communication repertoire (Fig. 3, Fig. S8). This observation indicates that poorly communicating species are not limited by structural barriers such as wing shape or olfactory capacity. Instead this suggests that, similar to local dialects in bird songs, Drosophila species-specific cues can be learned simply by repeated exposure to the “dialect”. We observed “dialect learning” in two different *D. ananassae* strain isolates, and two additional sister species (Fig. 3 B-E, Fig. S 8 A-G), indicating that dialect learning is likely to be a widespread phenomenon in Drosophila. Interestingly, some distantly related species that were unable to communicate with *D. melanogaster* (i.e. *D. willistoni*, *D. equinoxialis*) acquired the ability to partially communicate following a cohabitation-training period (Fig. 3 F-I, Fig. S 8 H-I). This was not the case for very distantly related species (i.e. *D. virilis*, *D. mojavensis*), which showed no ability to communicate with *D. melanogaster* even after training (Fig. 3 J-K, Fig. S 8 J-M). We also tested a transgenic *D. melanogaster*, to see if it was capable of teaching and dialect learning, and find such flies can teach their dialect to and learn the dialect from *D. ananassae* (Fig. S 8 N-O).

Additionally, we tested whether *D. ananassae* communication could benefit from cohabitation-training with species other than *D. melanogaster*. We find efficient communication between *D. simulans* (Fig. S 9 A-B), *D. equinoxialis* (Fig. S 9 C-D), and *D. mojavensis* (Fig. S 9 E-F) with *D. ananassae* following a cohabitation-training period. In contrast to the *D. melanogaster* results, we find communication with more distantly related species is altered after dialect training. *D. mojavensis and D. virilis* are able to dialect-learn following a cohabitation-training period with *D. ananassae*. With these species, in the untrained states we observe no ability to communicate (Fig. S 7 I-L), but find a partial communication phenotype following cohabitation (Fig. S 9 G-J). Therefore, *D. virilis* and *D. mojavensis*, although capable of interspecies communication and dialect learning, cannot learn the *D. melanogaster* dialect, but can learn *D. ananassae* dialect. These results suggest that some inter-species communication barriers do exist while others can be overcome by a period of dialect training and cohabitation.

Given our observation that two species can learn dialects following a cohabitation-training period, we wondered whether having more species present during the dialect training period influences dialect learning. In nature, flies are exposed to many different species of Drosophila. Given this, we hypothesized that neuronal plasticity exists in the fly brain to allow flies to learn multiple dialects from a given training period that includes multiple species as inputs. To probe this question, *D. melanogaster* were cohabitated with species capable of only partial communication or no communication in the untrained state, but show efficient and partial communication after dialect training (*i.e. D. ananassae* and *D. willistoni*, respectively). These three species were cohabitated for one week in a single container. We then used the trained *D. melanogaster* as students to *D. ananassae* and *D. willistoni* teachers (Fig. 4 A). We find that trained *D. melanogaster* are able to efficiently communicate with *D. ananassae* and partially communicate with *D. willistoni* (Fig. 4. B-C). These results mirror assays where these species were individually trained (Fig. 3 B-C, F-G), suggesting that flies can simultaneously make use of multiple inputs from multiple species and be able to learn and remember each unique dialect they encounter. Additionally, we tested *D. ananassae* and *D. willistoni* as students that were cohabitated with *D. melanogaster*. We find that *D. ananassae* can communicate efficiently with *D. melanogaster* and *D. willistoni* (Fig. 4 D-E), and that *D. willistoni* can partially communicate with *D. melanogaster* and effectively communicate with *D. ananassae* (Fig. 4 F-G). These data also mirror individual training (Fig. 3 B-C, F-G, Fig. S 9 E-F). Collectively, these data demonstrate that a fly can have vast communication repertoires consisting of multiple dialects that it acquires.

**Figure 4.**
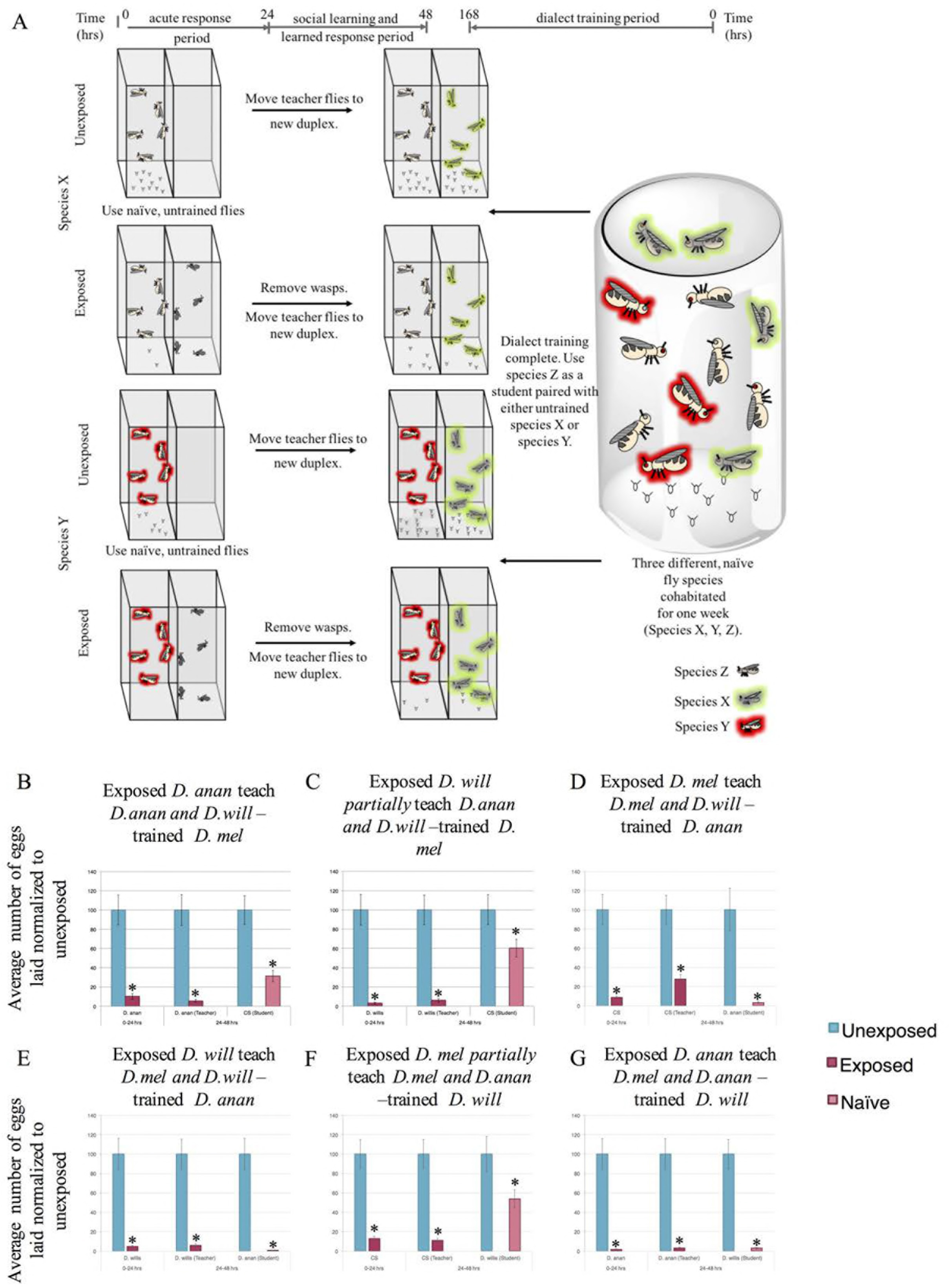
Flies can learn multiple dialects. (A) Experimental design of dialect training for flies that are used as students using multiple three unique species Percentage of eggs laid by exposed flies normalized to eggs laid by unexposed flies is shown. Communication between *D. melanogaster* students trained by *D. ananassae* and *D. willistoni*, shows that *D. melanogaster* learn each species dialect even in the presence of more than one species (B, C). Communication between *D. ananassae* students trained by *D. melanogaster* and *D. willistoni*, shows that *D. ananassae* learn each species dialect even in the presence of more than one species (D, E). Communication between *D. willistoni* students trained by *D. melanogaster* and *D. ananassae*, shows that *D. willistoni* learn each species dialect even in the presence of more than one species (F, G). Error bars represent standard error (n =12 biological replicates) (*p < 0.05).

### DIALECT LEARNING INPUTS

In order to better understand dialect learning, we tested the roles of sensory cues and genetic factors during the dialect-learning period. We measured dialect learning by quantifying improvement in interspecies partial communication between *D. melanogaster* and *D. ananassae* that normally exhibit efficient communication only after cohabitation. Given that in *D. melanogaster*, and in other species tested, we found visual cues to be sufficient for the teacher-student dynamic (Fig. 1) (17), we asked if visual cues are sufficient and/or necessary for dialect learning. We approached this question by performing the dialect training in the fly duplex, such that the two species could only see each other (Fig. 5 A), or by performing the training in the dark, so that the two species could physically interact, but lacked visual cues (Fig. S 10 A). We find that visual cues alone are not sufficient (Fig. 5 B-C), but are necessary (Fig. S 10 B-C) for dialect learning. The observation that visual cues are necessary but not sufficient makes the dialect learning phenomena fundamentally different from the teacher-student dynamic that requires only visual cues (17). Furthermore, we wondered if seeing another species altered the behavior of a fly to facilitate dialect learning. Blind *D. melanogaster ninaB* mutants do not function as students. Surprisingly, *D. ananassae* cohabitated with blind *D. melanogaster* do not learn the *D. melanogaster* dialect (Fig. S 10 D-E). We also performed cohabitation training under two different, monochromatic light sources, and this resulted in only a partial communication between *D. melanogaster* and *D. ananassae*, (Fig. 5 D-E, Fig. S 10 F-G). To exclude the possibility of a dimmer light source inhibiting dialect training under monochromatic settings, we repeated cohabitation-dialect-training in a full spectrum, lower light intensity setting, and found both species were able to learn the dialect (Fig. S 10 H-I). Thus, full spectrum light is essential in dialect learning. Importantly, the observation that blind *D. melanogaster* do not allow wild-type *D. ananassae* to dialect-learn suggests that species must see each other in order to alter their behavioral/chemical outputs required to facilitate dialect-learning.

**Figure 5.**
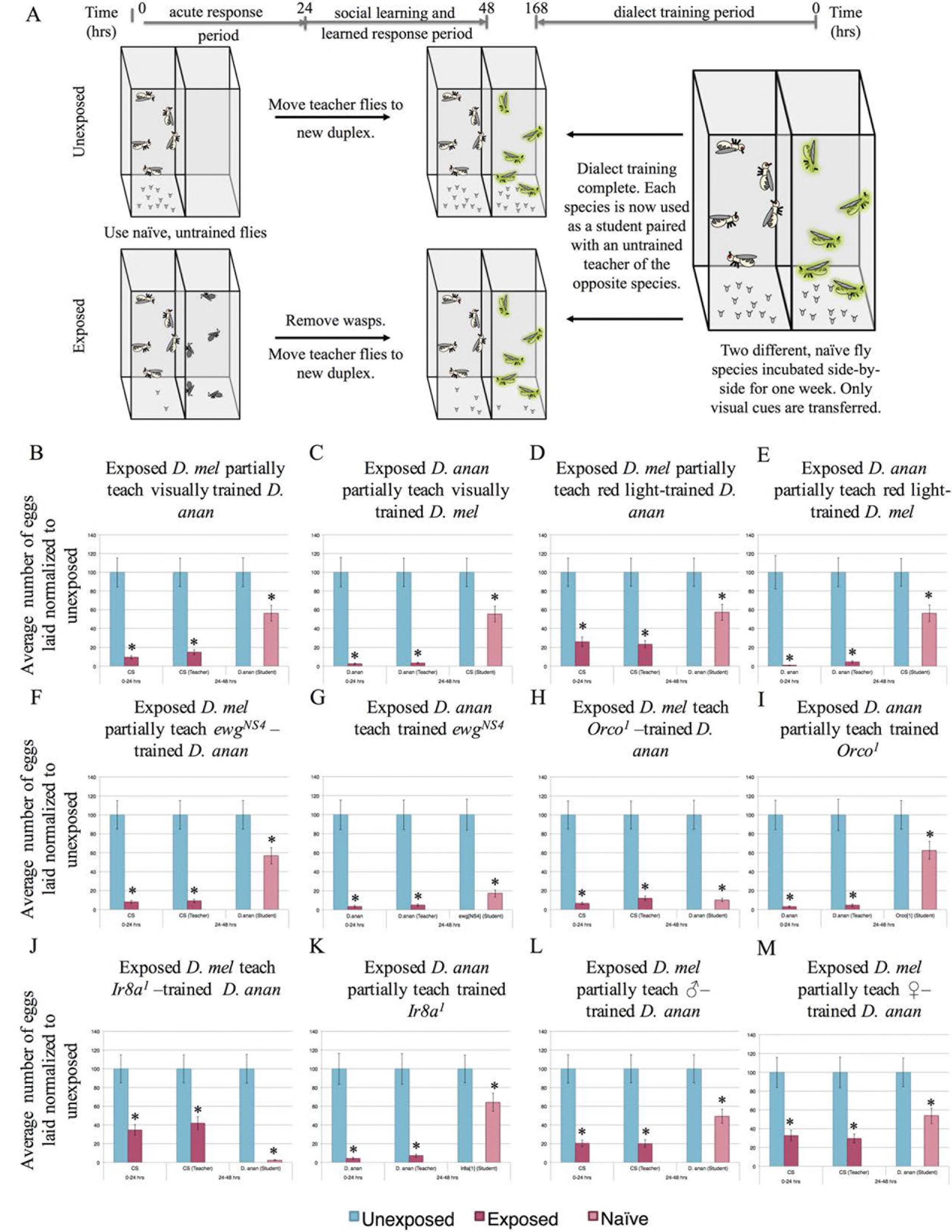
Dialect training requires multiple sensory inputs. (A) Experimental design of dialect training for flies that are used as students using only visual cues (panels B,C). Flies only see each other through the duplex, with no direct interaction. Two species are co-incubated for one week prior to being used as students. Percentage of eggs laid by exposed flies normalized to eggs laid by unexposed flies is shown. Communication between trained students *D. melanogaster* and *D. ananassae* with training through visual cues only, shows that visual cues are not sufficient (B, C). Communication between trained students *D. melanogaster* and *D. ananassae* with training through monochromatic, red light only, shows a lack of dialect training (D, E). Communication between trained students *ewg*^*NS4*^, mutant flies, and *D. ananassae* shows that moving wings are necessary (F, G). Communication between trained students *Orco*^*1*^ and *D. ananassae* shows that olfactory cues are necessary (H, I). Communication between trained students *Ir8a*^*1*^ and *D. ananassae* shows that *Ir8a* is a necessary receptor (J, K). Communication between trained students *D. melanogaster* and *D. ananassae* with training by male *D. melanogaster* only or by female *D. melanogaster* only, is not sufficient for dialect training (L,M). Error bars represent standard error (n = 12 biological replicates) (*p < 0.05).

Wing movement was shown to be required for teacher flies to instruct students in the teacher-student dynamic (17), raising the possibility that wing movement was also important for dialect learning. Therefore, we tested flies mutant in the erect wing gene (*ewg*), which impairs wing movement while maintaining morphologically normal wings. The allele tested has wild-type EWG protein expression in the nervous system, thus is only deficient in its non-neuronal functions, such as flight muscles (48). We find that *D. ananassae* cannot dialect learn from *ewg*^*NS4*^ flies (Fig. 5 F), although *ewg*^*NS4*^ mutants have no dialect learning impairment (Fig. 5 G). This suggests that dialect learning by *D. ananassae* requires *D. melanogaster* to have mobile wings.

To test if olfactory cues play a role in dialect learning, we utilized *D. melanogaster* mutants defective in chemosensory signaling. The majority of olfactory receptors require a co-receptor for wild-type function, including *Orco* (Or83b) for odorant receptors (49) and *Ir8a* or *Ir25a* for ionotropic receptors (50). Ir8a olfactory sensory neurons (OSNs) primarily detect acids and Ir25a OSNs detect amines, allowing us to probe specificity of detection. We find that *D. ananassae* are able to learn dialect from *Orco*^*1*^, *Ir8a*^*1*^, *Ir25a*^*2*^, single and *Ir8a*^*1*^;*Ir25a*^*2*^;*Orco*^*1*^ triple mutants and RNAi expressing *D. melanogaster* targeting each of these gene products (Fig. 4 H,J, Fig S 9 A-L). By contrast only *Ir25a*^*2*^ mutant and RNAi knockdown *D. melanogaster* were able to learn the *D. ananassae* dialect (Fig. 5 I,K, Fig S 11 A-L). These data demonstrate that Orco- and Ir8a-mediated olfactory inputs are required for dialect-learning. This further suggests that multiple olfactory cues play important roles in the dialect-learning period. We also find that *D. melanogaster* males and females are both required for dialect-training *D. ananassae* (Fig. 5 L-M, Fig. S 11 M-N) and that the length of the training period is also critical, as 24 hours is insufficient a period for dialect-learning (Fig. S 11 O-P). Thus, although the exact olfactory molecule(s) critical during a dialect-learning period are yet to be identified, we speculate that dialect-learning is a complex process requiring visual, olfactory and sex specific cues.

To examine the possibility that dialect training involves active learning mediated by neurons of the mushroom body, we utilized the GAL4 Gene-Switch system to transiently express a transgene specifically in the mushroom body (MB). Using the GAL4 Gene-Switch ligand system, RU486 (51) activates the GAL4 transcription factor, while administration of the vehicle (methanol) does not (51). RU486 was administered during the cohabitation period (or methanol for control), but not when flies were used as students post-dialect training (Fig. 6 A). Feeding of RU486 to the MB switch driver line does not impair dialect learning (Fig. S 12 A). We expressed the Tetanus toxin light chain (UAS-TeTx) specifically in the MB of *D. melanogaster* (to inhibit synaptic transmission during dialect training). We find that *D. ananassae* are able to learn the dialect of these MB inhibited flies (Fig. 6 B). However, *D. melanogaster* in which MB synaptic transmission is inhibited during the training period are unable to learn the *D. ananassae* dialect (Fig. 6 C). Control methanol only conditions (i.e. no RU486 ligand) with flies of identical genotypes do not show this defect (Fig. S 12 B). These data collectively indicate that visual and olfactory cues are required and possibly relayed to the MB, either directly or indirectly, to facilitate dialect learning. By contrast MB function does not appear to be important for *D. melanogaster* behavior(s) that enable *D. ananassae* to learn a dialect (Fig. S 6 B). Consistent with this idea, although *Orb2*^*ΔQ*^ mutants cannot function as students (Fig. 6 E) (17), *D. ananassae* nevertheless learns the *D. melanogaster* dialect from *Orb2*^*ΔQ*^ mutants (Fig. 6 D).

**Figure 6.**
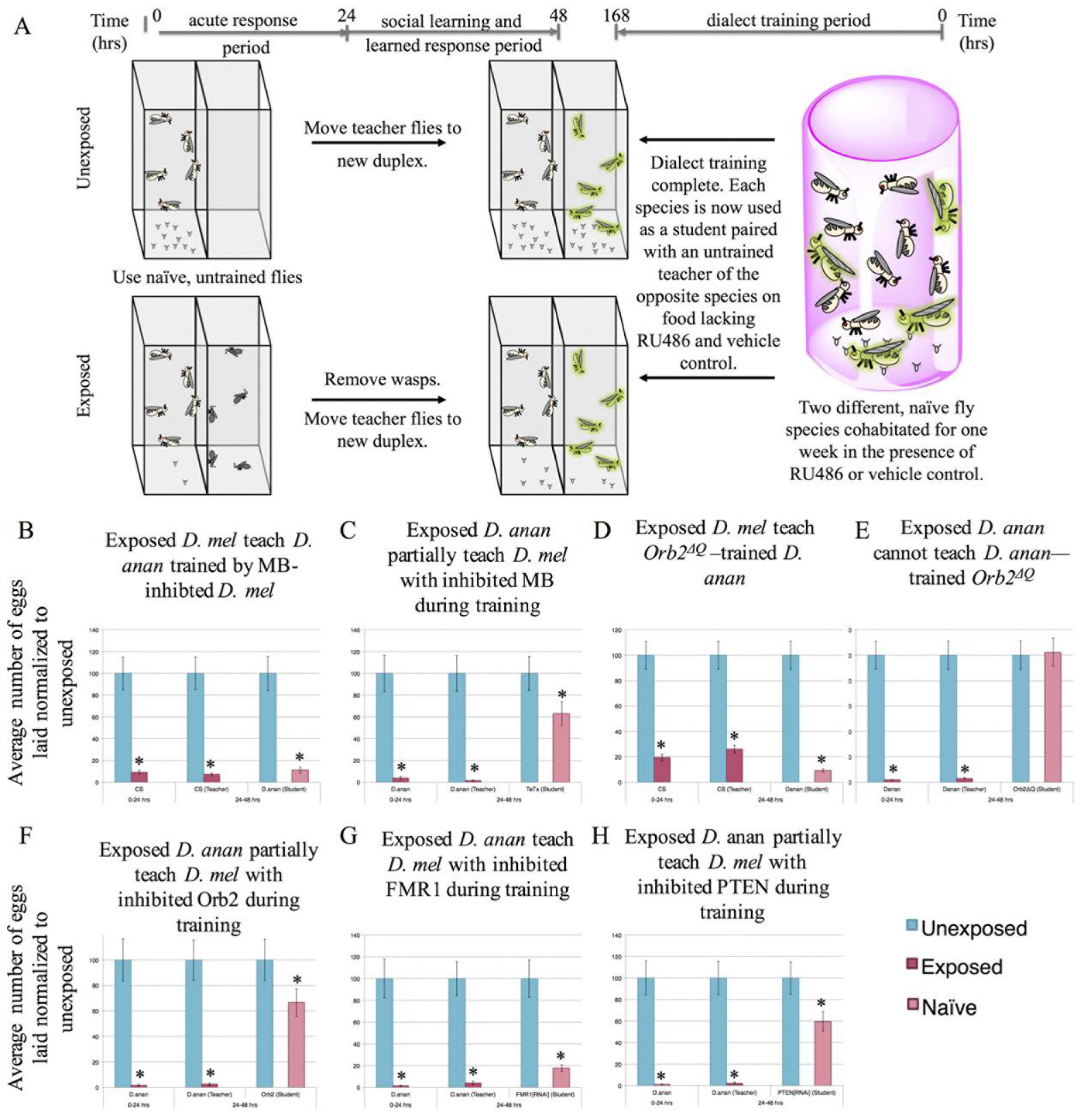
Genetic perturbations reveal a critical role of the mushroom body and memory proteins for dialect learning. (A) Experimental design of dialect training for flies being fed RU486 or methanol that are used as students. Both species are fed either RU486 or methanol during dialect training. Two species are co-incubated for one week prior to being used as students for naive, untrained teacher flies of the opposite species. Standard Drosophila media is used once the training period is over. Percentage of eggs laid by exposed flies normalized to eggs laid by unexposed flies is shown. Communication between trained students *D. melanogaster* and *D. ananassae* trained by flies expressing tetanus toxin (UAS-TeTx) in the mushroom body (MB) shows that the MB serves a critical role during the training period. *D. ananassae* learn from *D. melanogaster* with an inhibited MB, demonstrating that a functional MB is not needed to confer information during the training period (B, C). Communication between trained students *Orb2*^*ΔQ*^ and *D. ananassae* shows that Orb2 is required in students, but is dispensable for teachers to *D. ananassae* (D, E). Communication between *D. ananassae* and students co-incubated with *D. ananassae* that have RNAi-mediated Orb2 knockdown in the MB through RU486 feeding shows that the MB requires Orb2 during the training period (F). Communication between *D. ananassae* and students coincubated with *D. ananassae* that have RNAi-mediated FMR1 knockdown (strain #24944) in the MB through RU486 feeding shows that FMR1 is not required in the MB during the training period (G). Communication between *D. ananassae* and students co-incubated with *D. ananassae* that have RNAi-mediated PTEN knockdown in the MB through RU486 feeding shows that PTEN is required in the MB during the training period (H). Error bars represent standard error (n = 12 biological replicates) (*p < 0.05).

Because MB function is necessary for dialect learning during dialect training, we tested the long-term memory proteins Orb2, FMR1, and phosphatase and tensin homolog (PTEN) (52,53) that are known to be required in the MB for memory formation. PTEN has been implicated in murine social learning models, though it has not been tested in a social learning assay in Drosophila (54). We used the MB Gene-Switch to knockdown expression only during the cohabitation period, after which expression was allowed to resume. *D. ananassae* learn the dialect of each of these three knockdown lines, again suggesting that MB mediated processes in *D. melanogaster* are not necessary for *D. ananassae* dialect-training (Fig. S 12 C-G). However, under these conditions we find that functional Orb2 and PTEN are required for dialect learning in *D. melanogaster*, but FMR1 is dispensable (Fig. 6 F-H). Orb2 and FMR1 were previously shown to be important in the teacher-student transmission of a wasp threat, and knockdown of either gene completely ablated students learning from teacher flies. In this case, partial communication between *D. ananassae* teachers and *D. melanogaster* students can occur because Orb2 and PTEN expression is restored after the dialect-training period, thus functioning as wild-type *D. melanogaster*. *D. melanogaster* flies having undergone knockdown of Orb2 and PTEN only during dialect-training are able to communicate with and function as students to wild-type *D. melanogaster* after the cohabitation period is completed, suggesting the partial communication phenotype observed with *D. ananassae* teachers is a result of gene knockdown during cohabitation and not a by-product of irreversible cellular damage or death caused by the RNAi treatments (Fig. S 12 H-L). Collectively, these data show critical gene products are required to function in the MB for dialect learning during the training period. Importantly, MB function and active learning are not necessary in *D. melanogaster* in order to in turn provide cues enabling dialect learning by a wild-type *D. ananassae* student.

## DISCUSSION

In this study, we present an evolutionarily conserved response to predatory wasps across the genus Drosophila, manifesting as oviposition depression coincident with an activated effector caspase, Dcp-1. We have shown that flies communicate a wasp threat through visual cues. Interspecies communication occurs to varying degrees, likely dependent on evolutionary relatedness. Closely related species, such as *D. melanogaster* and *D. simulans*, *D. ananassae* and *D. kikkawai*, and *D. mojavensis* and *D. virilis*, communicate as effectively as conspecifics. Species more distantly related to *D. melanogaster* exhibit only partial communication or lack the ability to confer predator information with *D. melanogaster*. When two species are only able to partially communicate, they can learn each other's dialect after a period of cohabitation, yielding inter-species communication enhanced to levels normally observed among conspecifics. Although dialect learning facilitates inter-species communication across broad evolutionary distances, the ability to learn a specific dialect is dependent on relatedness of the two species (Fig. 7 A). This observation of the role of phylogenetic distance influencing dialect learning is true in cases both utilizing *D. melanogaster* and *D. ananassae* in combination with other species tested (Fig. 7 A, Fig. S 13). The observation that different strains of the same species exhibit this partial communication that can then be enhanced by cohabitation, suggests that both social communication and dialect learning are innate behaviors conserved among all Drosophilids tested here (Fig. 7 A, Fig. S 13). Multiple strains of *D. melanogaster* reared in the laboratory for many decades exhibit this behavior, supporting the idea that this is an innate behavior. Thus, adult Drosophila neuronal plasticity allows for learning of dialects, but the specific dialect learned is dependent on social interactions specific to a communal environmental context that provides both visual and olfactory inputs. This same plasticity allows for the learning of multiple dialects in a given environment. It is remarkable that communal rearing of two species can enhance communication about a predator that is yet to be experienced by either species. Furthermore, dialect-learning does not trigger Dcp-1 activation and oviposition depression, suggesting that social communication about predator presence is different from social interactions that enable dialect-learning that later enhances predator presence communication.

**Figure 7.**
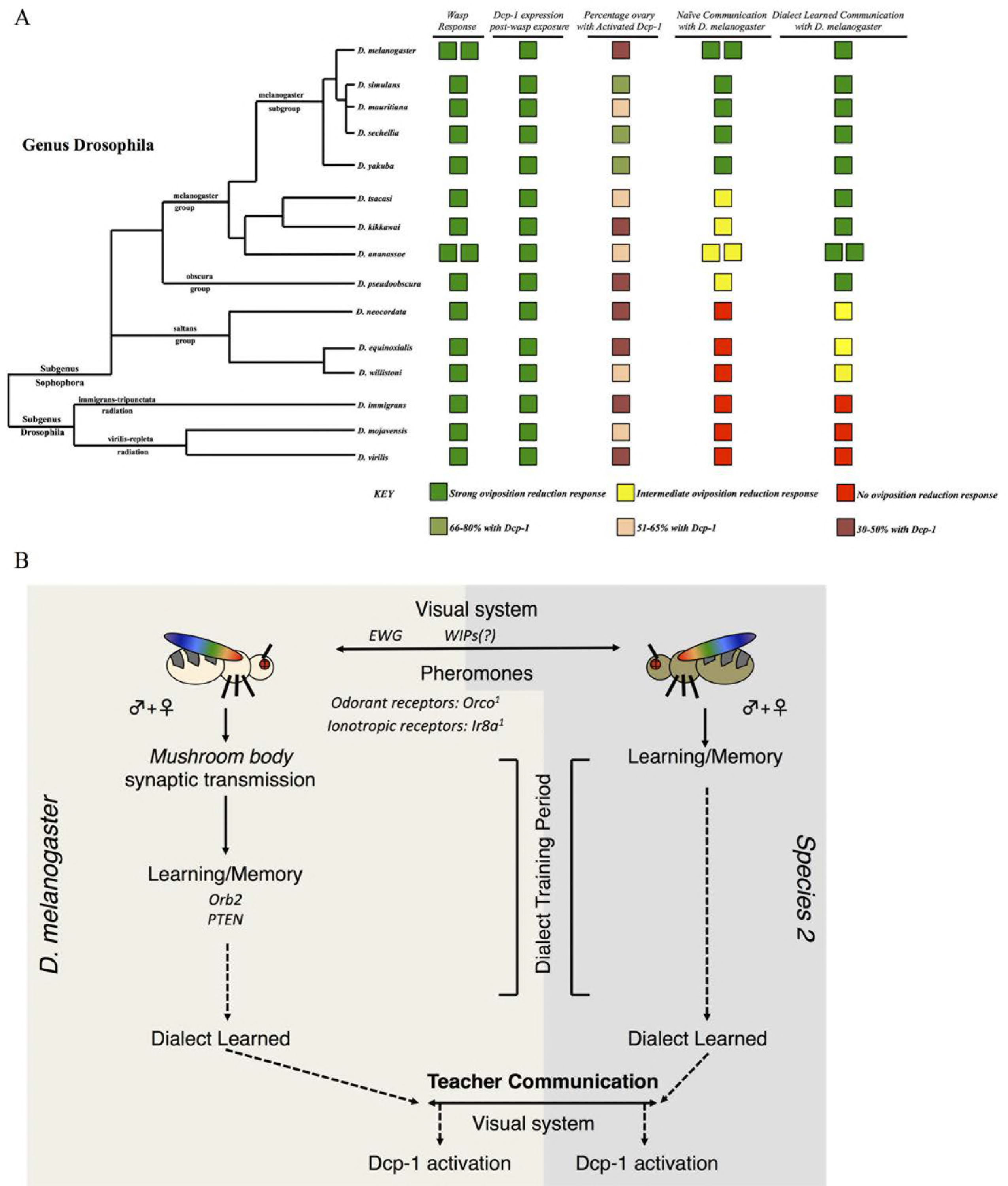
Phylogenetic summary of dialect learning and pathway model for interspecies social learning. Utilizing species across the genus Drosophila (A) demonstrates conservation of oviposition depression following wasp exposure, mediated by activated Dcp-1 to varying degrees and with varying expression patterns. The ability to communicate with *D. melanogaster* and the ability to demonstrate interspecies communication varies across the genus, with species closely related to *D. melanogaster* able to communicate without barriers. More distantly related species have difficulty communicating, though the barrier can be alleviated with dialect training. Finally, some species are too distantly related to communicate even after dialect training. Double boxes in a given row and column indicate multiple wild-type strains were tested. Interspecies communication is dependent on the presence of both male and female flies, the visual and olfactory systems, the mushroom body, and various long-term memory gene products (B). This model is based of the use of *D. melanogaster* and *D. ananasse*. Alleles tested in (B) are: Orco[1], Ir8a[1];Ir25a[2];Orco[1], Ir8a[1], Ir25a[2], ninaB[P315], Orb2AQ, ewg[NS4], Orb2[RNAi], PTEN[RNAi], FMR1[RNAi], and UAS-TeTx.

We propose dialect-learning to be a novel behavior requiring visual and olfactory inputs, perhaps integrated in and relayed through the MB, resulting in the ability to more efficiently receive information about a common predator. Without dialect learning, this information would otherwise be lost in translation or muddled, resulting in an inefficient behavioral response with significant survival disadvantages. Inhibiting synaptic transmission and knockdown of key learning and memory genes in the MB demonstrates that these inputs must be processed and consolidated in the MB, although input neuronal signaling is initiated from the visual and olfactory systems (Fig. 7 B). Given the need for multiple sensory inputs, dialect learning is fundamentally different from the previously described teacher-student paradigm, where visual cues are necessary and sufficient for information exchange (17). Additionally, we suggest that this study also points to previously unappreciated functions of the Drosophila MB in integrating information from multiple olfactory and visual inputs. Such cognitive plasticity that allows for dialect learning from many different species hints that adult behaviors could only emerge in a manner that is dependent on previous social experiences where relevant ecological pressures are ever present and multiple species co-exist in nature. Thus, there is a real benefit to cognitive plasticity, where sharing of information directly, or by coincident bystanders, could result in behavioral immunity to pan-specific threats.

The specific information shared by different species during dialect learning is not known. This study, however, provides important clues as the complex suite of sensory systems and cues that may be required for efficient dialect learning. Visual sensory input is critical in dialect learning and it is intriguing that both wing movement and full spectrum light are essential. This observation raises the very interesting possibility that dialect learning may require wing interference patterns (WIPs) *via* wing movement in the presence of full spectrum light (55,56) (Fig. 7 B). WIPs are known to be produced by species-specific wing patterns and light diffraction abilities and in Drosophila are a source of information for making mate choice decisions (CITE). Given that visual, wing movement based cues are required for dialect learning, we speculate that in full spectrum light WIPs could facilitate dialect learning in closely related species, while more divergent WIPs could also prohibit distantly related species from communicating at all.

We have presented an example of how inter-species social communication and dialect learning in Drosophila can lead to changes in germline physiology and reproductive behavior. What other ethological behaviors are modulated by MB functions and social interactions typically not revealed in laboratory monocultures? We suggest that the Drosophila MB may integrate a myriad of social and environmental cues in order to produce ethologically relevant behavior that is responsive and useful to local environmental conditions.

## Acknowledgements

We thank Yashi Ahmed, Greg Roman, Greg Suh, FlyBase, the Bloomington Drosophila Stock Center, and the Drosophila Species Stock Center (DSSC) at the University of California, San Diego, for stocks. We thank Theresa Reimels, Erin Kelleher, and Mani Ramswami for helpful comments on the manuscript. We also thank the Dartmouth Department of Biological Sciences Light Microscopy Facility for technical assistance. We acknowledge grants from Geisel School of Medicine at Dartmouth, the National Institute of Health Pioneer grant:1DP1MH110234 (GB), and the Defense Advanced Research Projects Agency, grant:HR0011-15-1-0002 (GB).

## SUPPLEMENTARY FIGURE LEGENDS

**Supplementary Figure 1.**
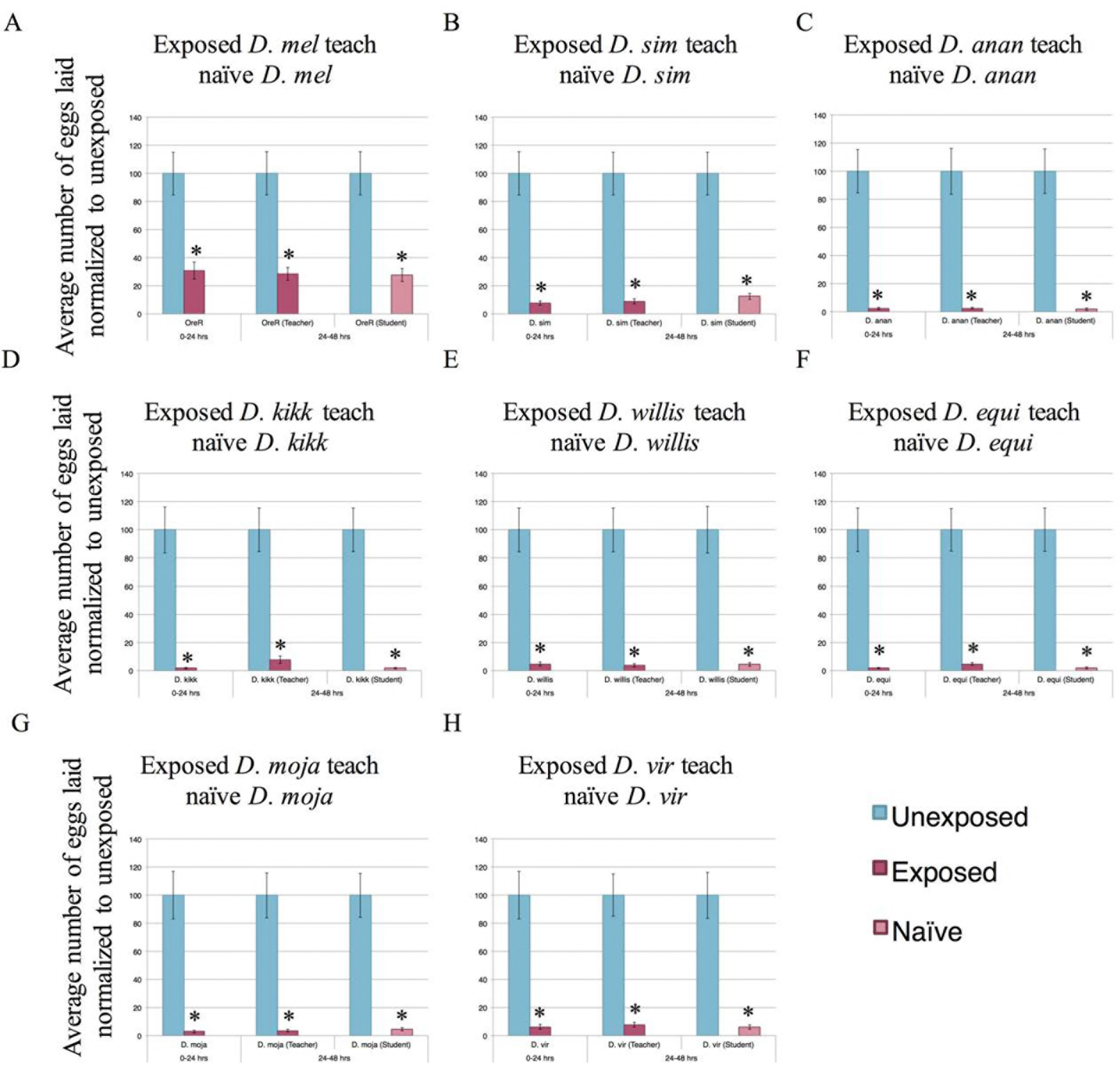
Intra-species communication is present across the genus Drosophila. Percentage of eggs laid by exposed flies normalized to eggs laid by unexposed flies is shown. Species shown are (A) *D. melanogaster* (Oregon-R), (B) *D. simulans*, (C) *D. ananassae*, (D) *D. kikkawai*, (E) *D. willistoni*, (F) *D. equinoxialis*, (G) *D. mojavensis*, and (H) *D. virilis*. Error bars represent standard error (n = 12 biological replicates) (*p < 0.05).

**Supplementary Figure 2.**
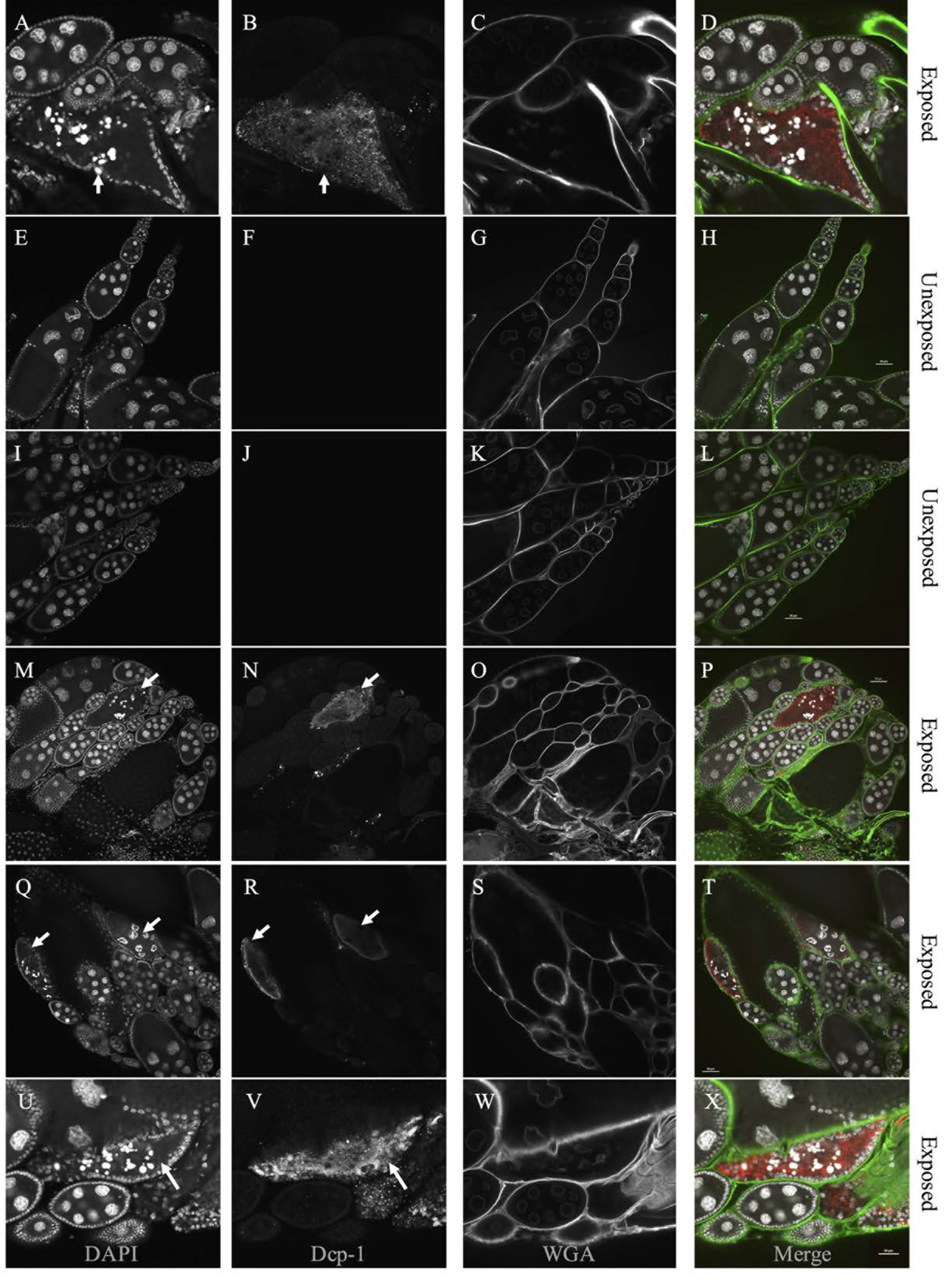
Activated Dcp-1 is indicative of apoptotic events in *D. melanogaster*. Magnified images from Figure 1 (H-K) showing apoptotic egg chamber displaying activated caspase. DAPI (A), activated Dcp-1 (B), WGA (C) and merge are shown (D). Additional representative ovaries of unexposed and wasp-exposed *D. melanogaster* are shown. DAPI (E, I, M, Q, U), activated Dcp-1 (F, J, N, R,V), WGA (G, K, O, S, W), and the merged images (H, L, P, T, X) are shown. Arrows denote apoptotic egg chambers.

**Supplementary Figure 3.**
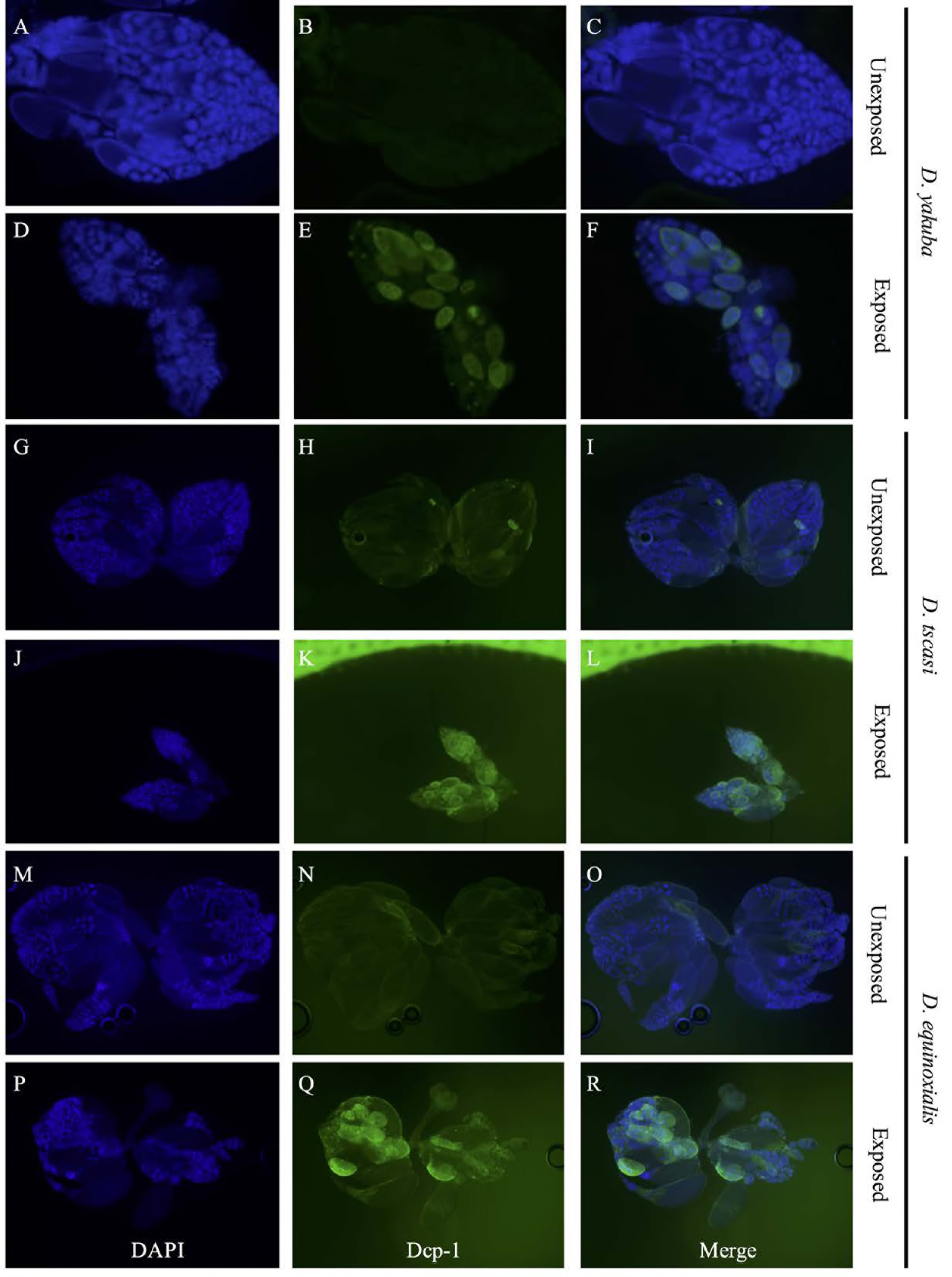
Increases in activated 763 caspase are observed in the ovary across the genus Drosophila following wasp exposure. Representative images of unexposed and wasp-exposed ovaries stained for activated Dcp-1 for *D. yakuba* (A-F), *D. tsacasi* (G-L), and *D. equinoxialis* (M-R). DAPI, Dcp-1, and the merged images are shown. The broad range of staining patterns observed in these species is representative of other species tested.

**Supplementary Figure 4.**
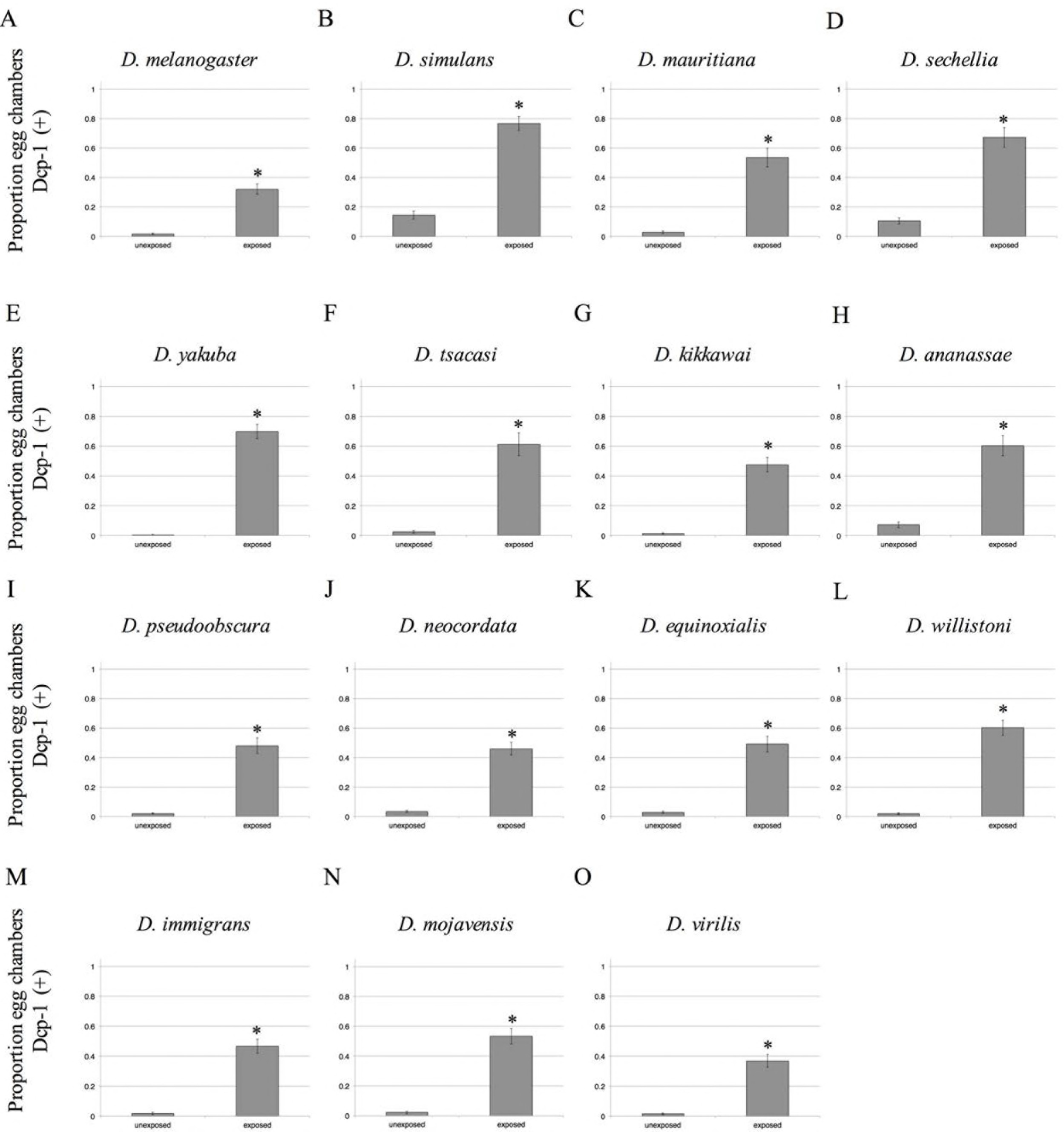
Increases in activated caspase are quantified in the ovary across the genus Drosophila following wasp exposure. Proportion of egg chambers with Dcp-1 signal shown for (A) *D. melanogaster*, (B) *D. simulans*, (C) *D. mauritiana*, (D) *D. sechellia*, (E) *D. yakuba*, (F) *D. tsacasi*, (G) *D. kikkawai*, (H) *D. ananassae*, (I) *D. pseudoobscura*, (J) *D. neocordata*, (K) *D. equinoxialis*, (L) *D. willistoni*, (M) *D. immigrans*, (N) *D. mojavensis*, and (O) *D. virilis*. Error bars represent standard error (n = 36 ovaries) (*p < 0.05).

**Supplementary Figure 5.**
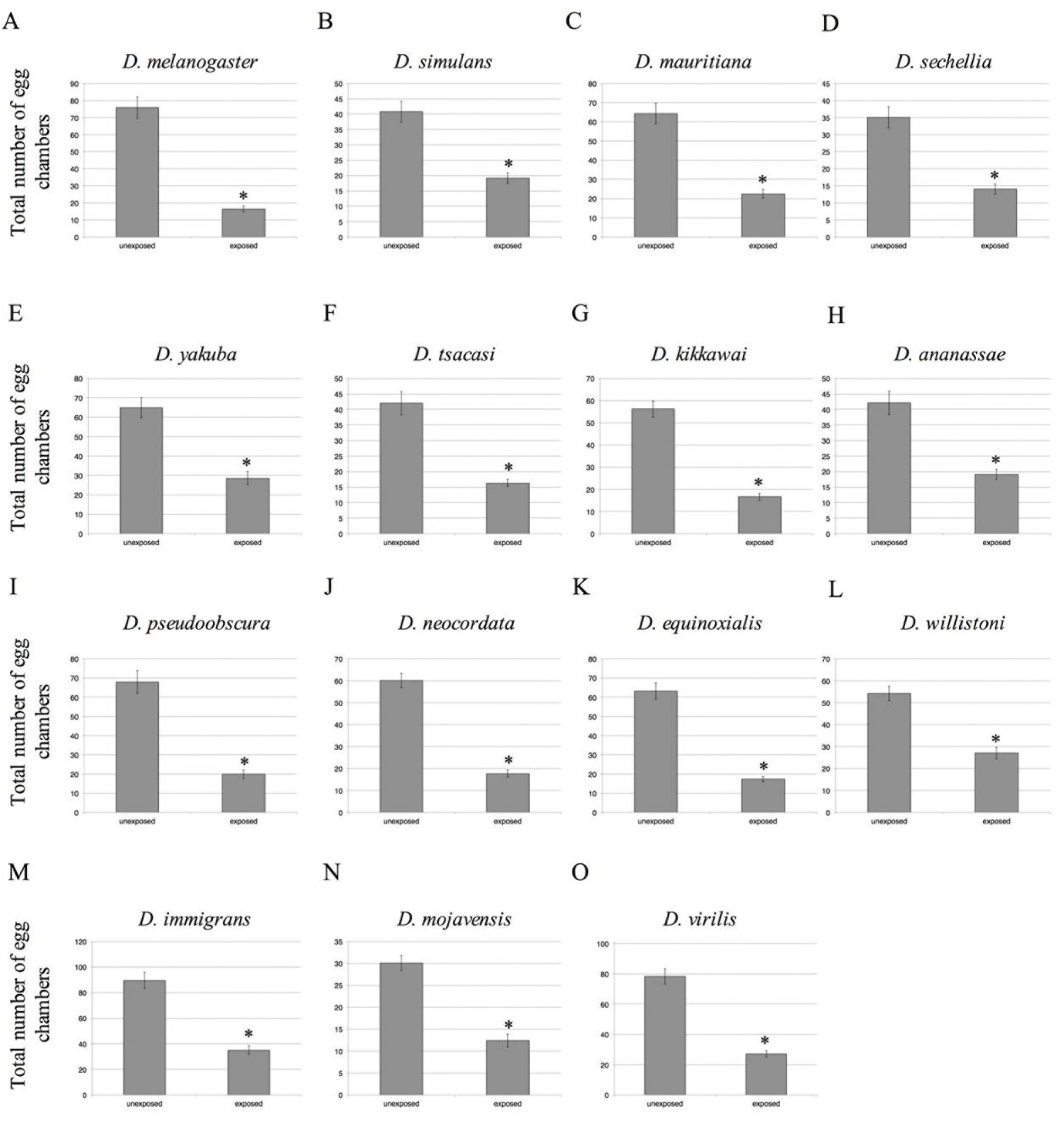
A decrease in egg chamber numbers are quantified in the ovary across the genus Drosophila following wasp exposure. Total number of egg chambers shown for (A) *D. melanogaster*, (B) *D. simulans*, (C) *D. mauritiana*, (D) *D. sechellia*, (E) *D. yakuba*, (F) *D. tsacasi*, (G) *D. kikkawai*, (H) *D. ananassae*, (I) *D. pseudoobscura*, (J) *D. neocordata*, (K) *D. equinoxialis*, (L) *D. willistoni*, (M) *D. immigrans*, (N) *D. mojavensis*, and (O) *D. virilis*. Error bars represent standard error (n = 36 ovaries) (*p < 0.05).

**Supplementary Figure 6.**
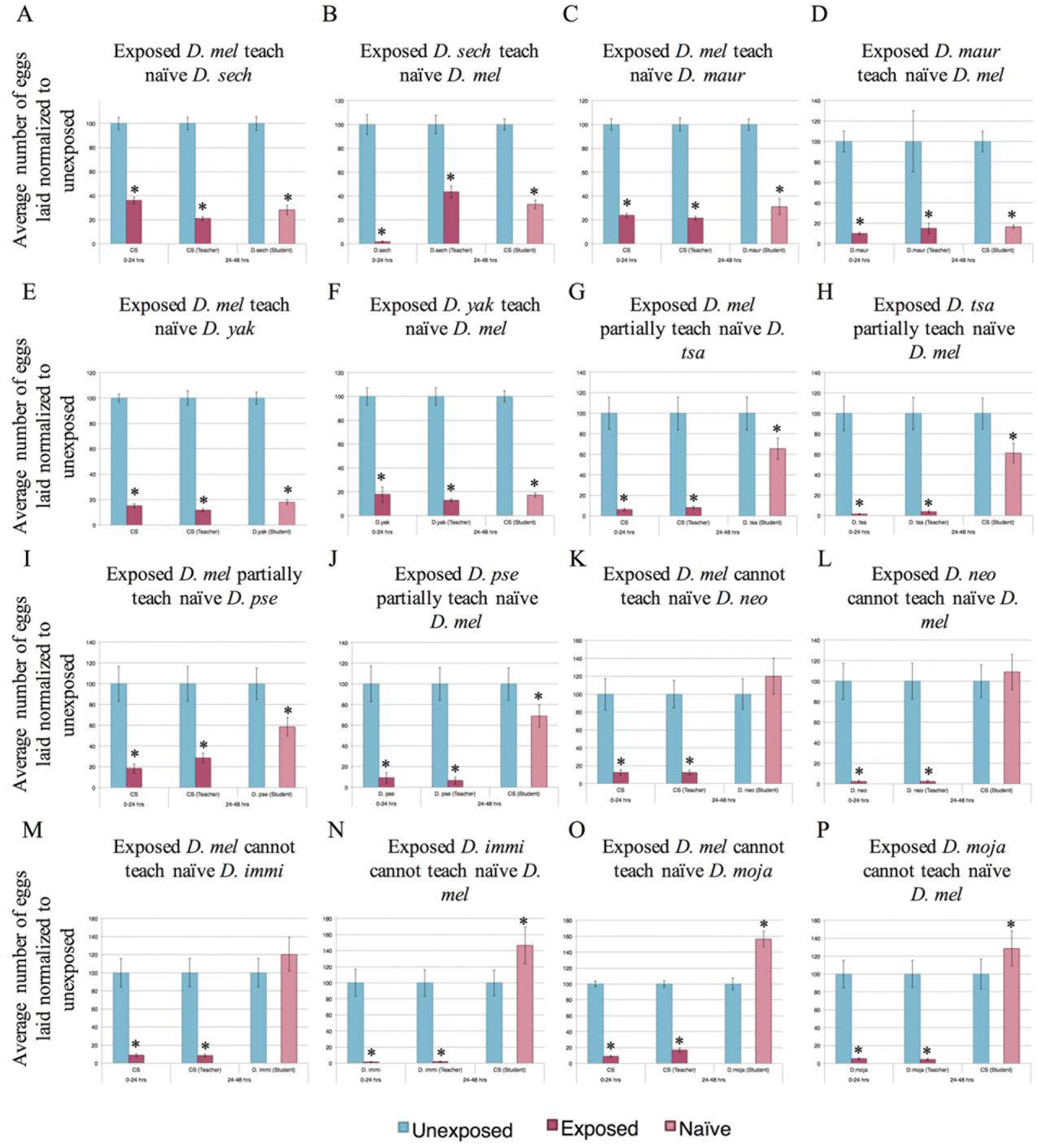
Interspecies communication of predator threats. Percentage of eggs laid by exposed flies normalized to eggs laid by unexposed flies is shown. Communication between *D. melanogaster* and: *D. sechellia* (A, B), *D. mauritianna* (C, D), *D. yakuba* (E, F), *D. tsacasi* (G, H), *D. pseudoobscura* (I, J), *D. neocordata* (K, L), *D. immigrans* (M, N), and *D. mojavensis* (O, P), shows varying communication abilities. Error bars represent standard error (n = 12 biological replicates) (*p < 0.05).

**Supplementary Figure 7.**
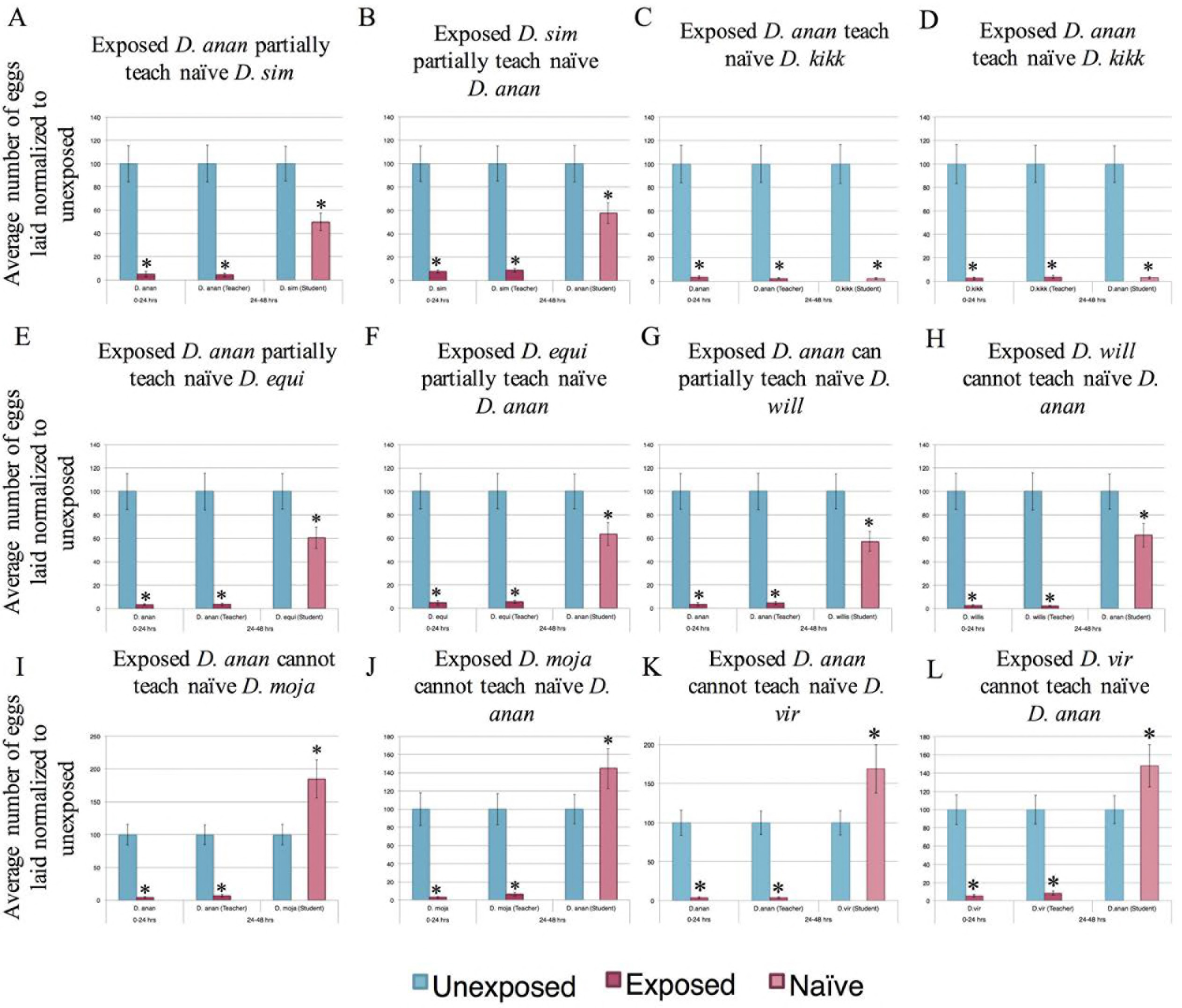
Interspecies communication of predator threats using *D. ananassae*. Percentage of eggs laid by exposed flies normalized to eggs laid by unexposed flies is shown. Communication between *D. ananassae* and: *D. simulans* (A, B), *D. kikkawai* (C, D), *D. equinoxialis* (E, F), *D. willistoni* (G, H), *D. mojavensis* (I, J), and *D. virilis* (K, L), shows varying communication abilities. Error bars represent standard error (n = 12 biological replicates) (*p < 0.05).

**Supplementary Figure 8.**
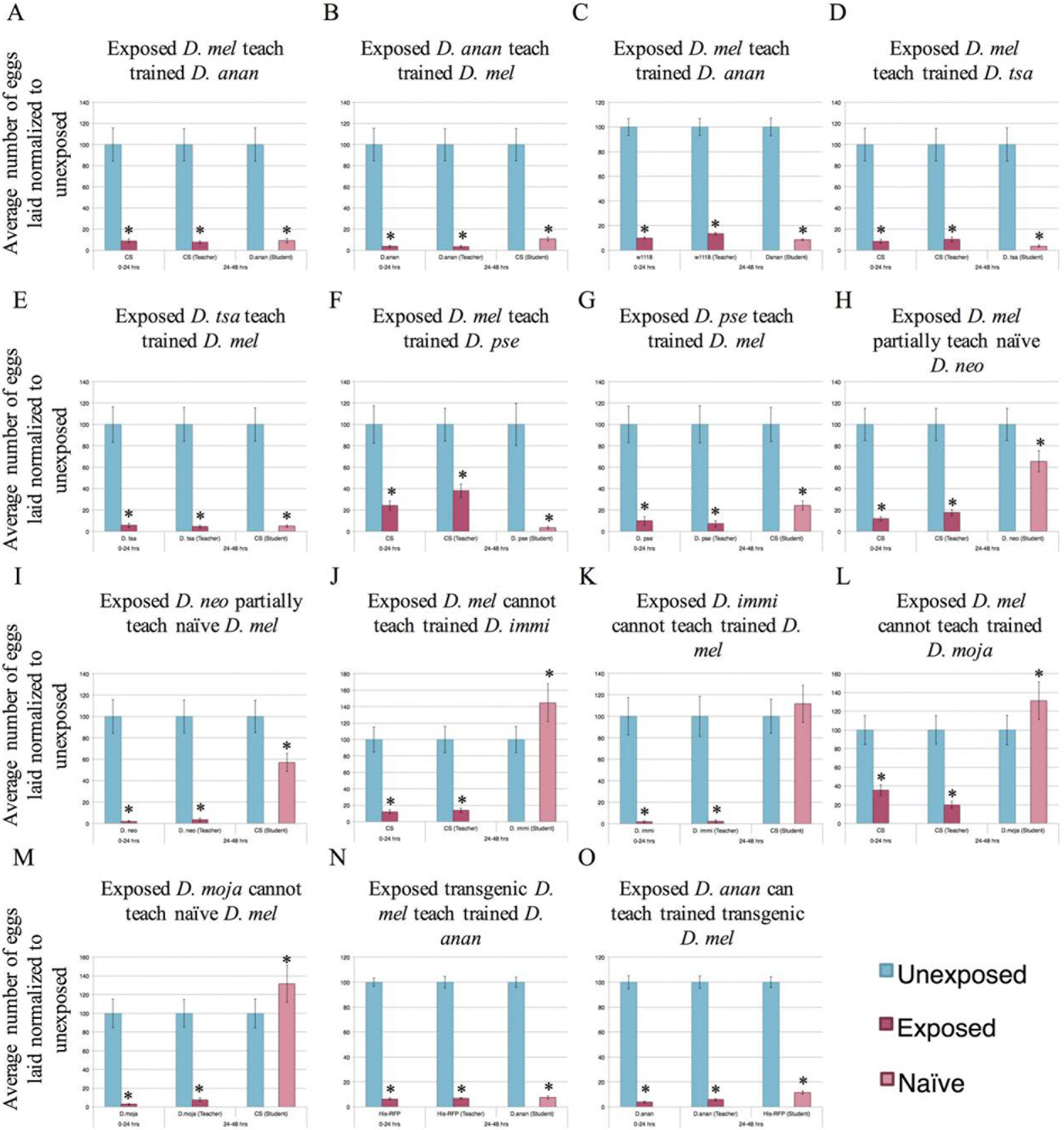
Cohabitation of additional species with D. melanogaster allows for interspecies communication. Percentage of eggs laid by exposed flies normalized to eggs laid by unexposed flies is shown. Communication between trained students *D. melanogaster* and: *D. ananassae* (second line) (A-C), *D. tsacasi* (D, E), *D. pseudoobscura* (F, G), *D. neocordata* (H, I), *D. immigrans* (J, K), and *D. mojavensis* (L, M). (C) An additional *D. melanogaster* line (*w*^*1118*^) learns from *w*^*1118*^ trained *D. ananassae*. Communication between *D. ananassae* and a transgenic *D. melanogaster* (Histone-RFP) occurs following training period (N, O). Error bars represent standard error (n = 12 biological replicates) (*p < 0.05).

**Supplementary Figure 9.**
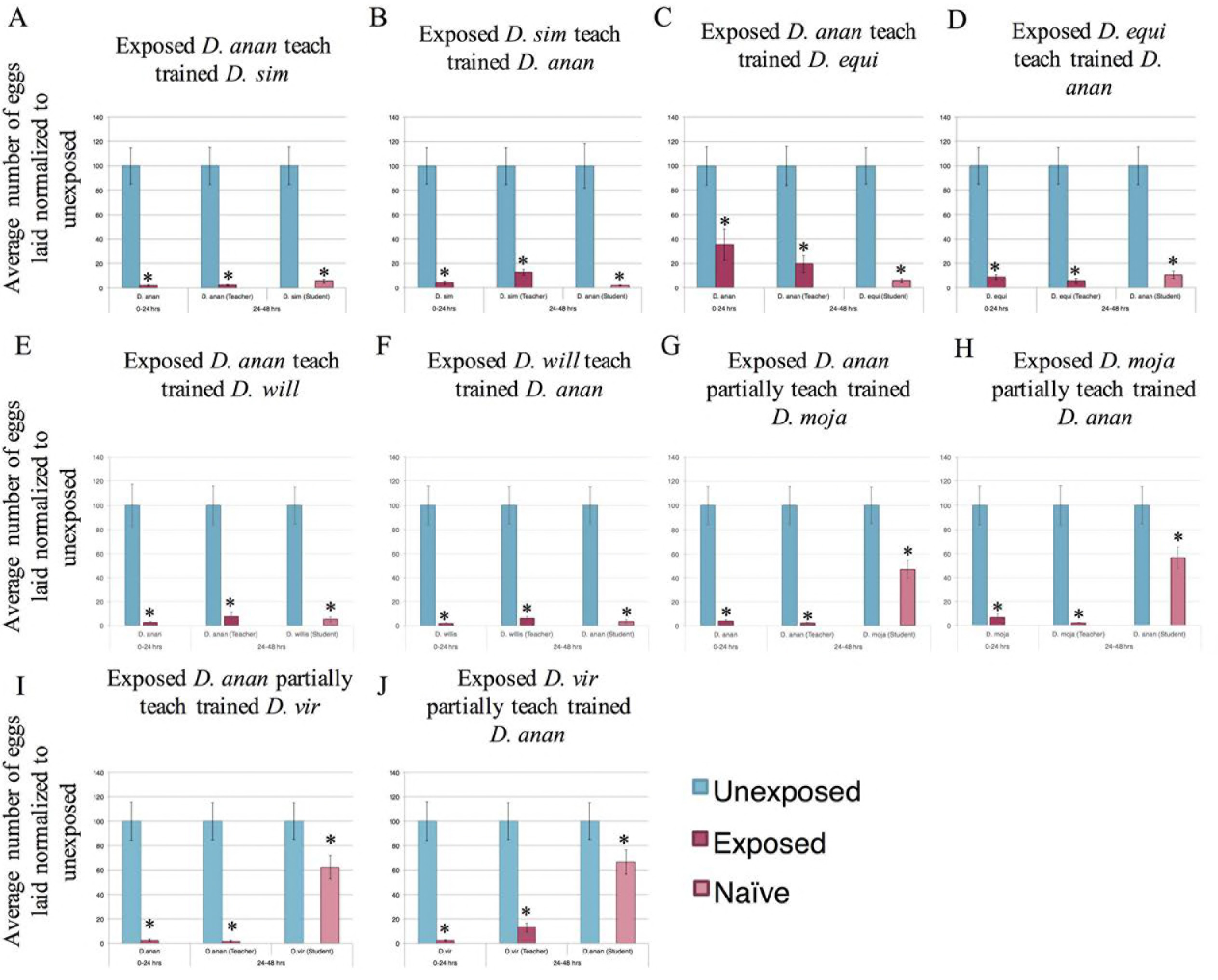
Cohabitation of additional species with D. ananassae allows for interspecies communication. Percentage of eggs laid by exposed flies normalized to eggs laid by unexposed flies is shown. Communication between trained students *D. ananassae* and: *D. simulans* (A,B), *D. equinoxialis* (C,D), *D. willistoni* (E,F), *D. mojavensis* (G,H), and *D. virilis* (I,J). Error bars represent standard error (n = 12 biological replicates) (*p < 0.05).

**Supplementary Figure 10.**
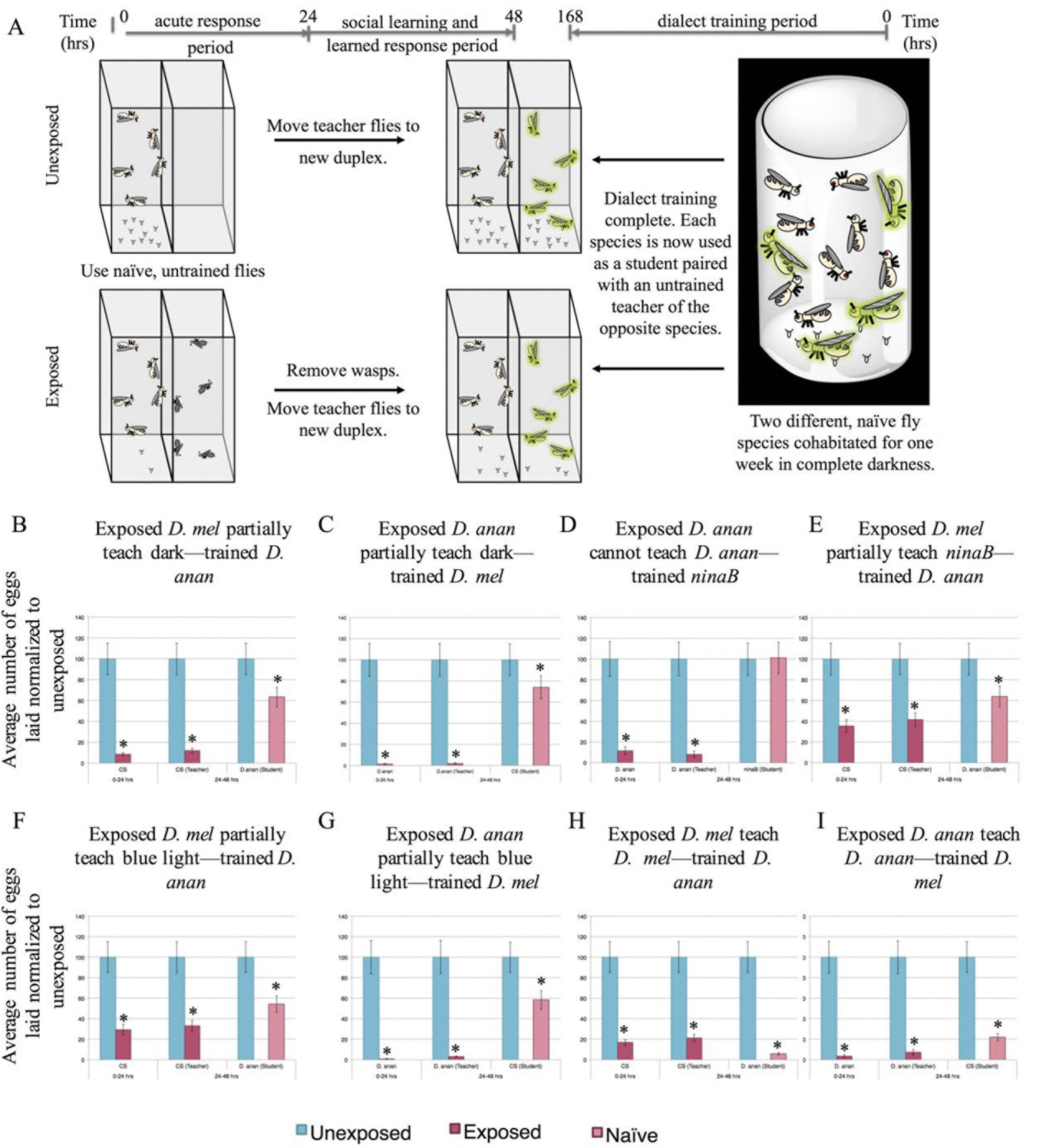
Additional evidence demonstrating that dialect training requires visual cues. (A) Experimental design of dialect training for flies that are used as students using no visual cues by running the dialect training period in the dark (B,C). Flies do not see each other, but still interact and innervate other sensory inputs. The two species are co-incubated for one week prior to being used as students for naive, untrained teacher flies of the opposite species. Percentage of eggs laid by exposed flies normalized to eggs laid by unexposed flies is shown. Communication between trained students *D. melanogaster* and *D. ananassae* with training involving no visual cues (dark-trained), shows that visual cues necessary for dialect learning (B, C). Communication between trained students *D. ananassae* and the mutant *ninaB* (D,E). Communication between trained students *D. melanogaster* and *D. ananassae*, with training in monochromatic blue light only, shows a lack of dialect training (F, G). Communication between trained students of *D. ananassae* and *D. melanogaster* at 4.0_8_ light intensity shows communication (H,I). Error bars represent standard error (n = 12 biological replicates) (*p < 0.05).

**Supplementary Figure 11.**
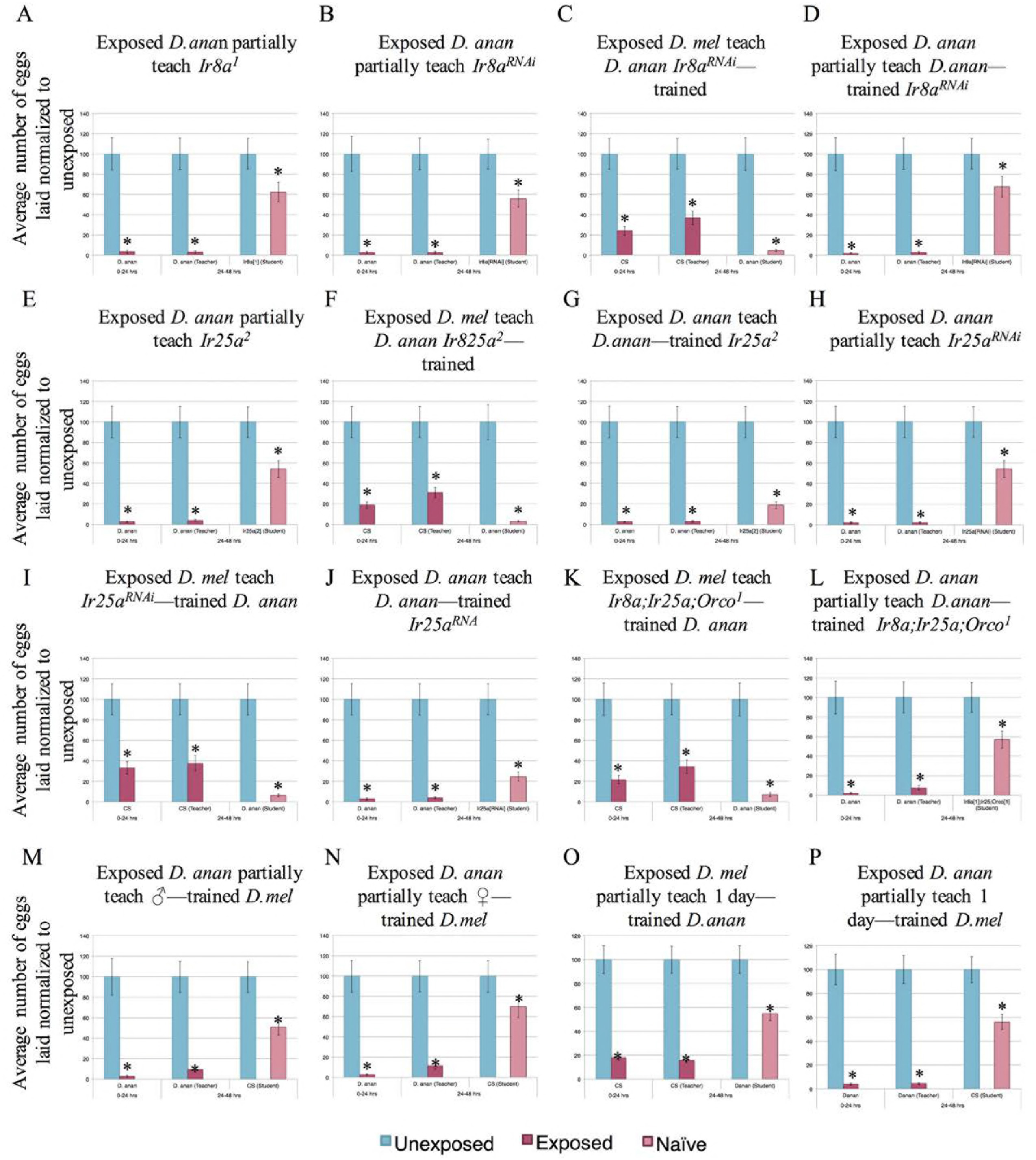
Further evidence demonstrating that dialect training requires multiple sensory inputs including olfactory cues and duration of training. Percentage of eggs laid by exposed flies normalized to eggs laid by unexposed flies is shown. Communication between naïve *D. ananassae* and *IrSa*^*1*^ mutant flies shows partial communication (A). Communication between naïve students of *IrSa* knockdown in Ir8a-expressing neurons and *D. ananassae* shows partial communication (B). Communication between trained students *IrSa*^*RNAl*^ knockdown in Ir8a expressing neurons and *D. ananassae* shows that IR8a receptor-mediated cues are necessary (C, D). Communication between naïve *Ir25a*^*2*^ mutants and *D. ananassae* shows partial communication (E). Communication between trained students *Ir25a*^*2*^ and *D. ananassae* shows communication suggesting that IR25a receptors are not required for dialect training (F,G). Communication between naïve *Ir25a* knockdown in Ir25a-expressing neurons and *D. ananassae* shows partial communication (H). Communication between trained students *Ir25a*^*mAi*^ knockdown in Ir25a-expressing neurons and *D. ananassae* shows communication suggesting that IR25a receptors are not required for dialect training (I, J). Communication between trained *IrSa*^*1*^*;Ir25a*^*2*^*;Orco*^*1*^ students and *D. ananassae* shows that olfactory and IR-receptor mediated cues are necessary (K, L). Communication between students *D. melanogaster* and *D. ananassae*, with training by males only or by females only, shows partial communication, suggesting that both male and female flies are required for dialect learning (M, N). Communication between trained students *D. melanogaster* and *D. ananassae*, with training for only one day, shows that 24 hours is not sufficient for dialect training (O, P). Error bars represent standard error (n = 12 biological replicates) (*p < 0.05).

**Supplementary Figure 12.**
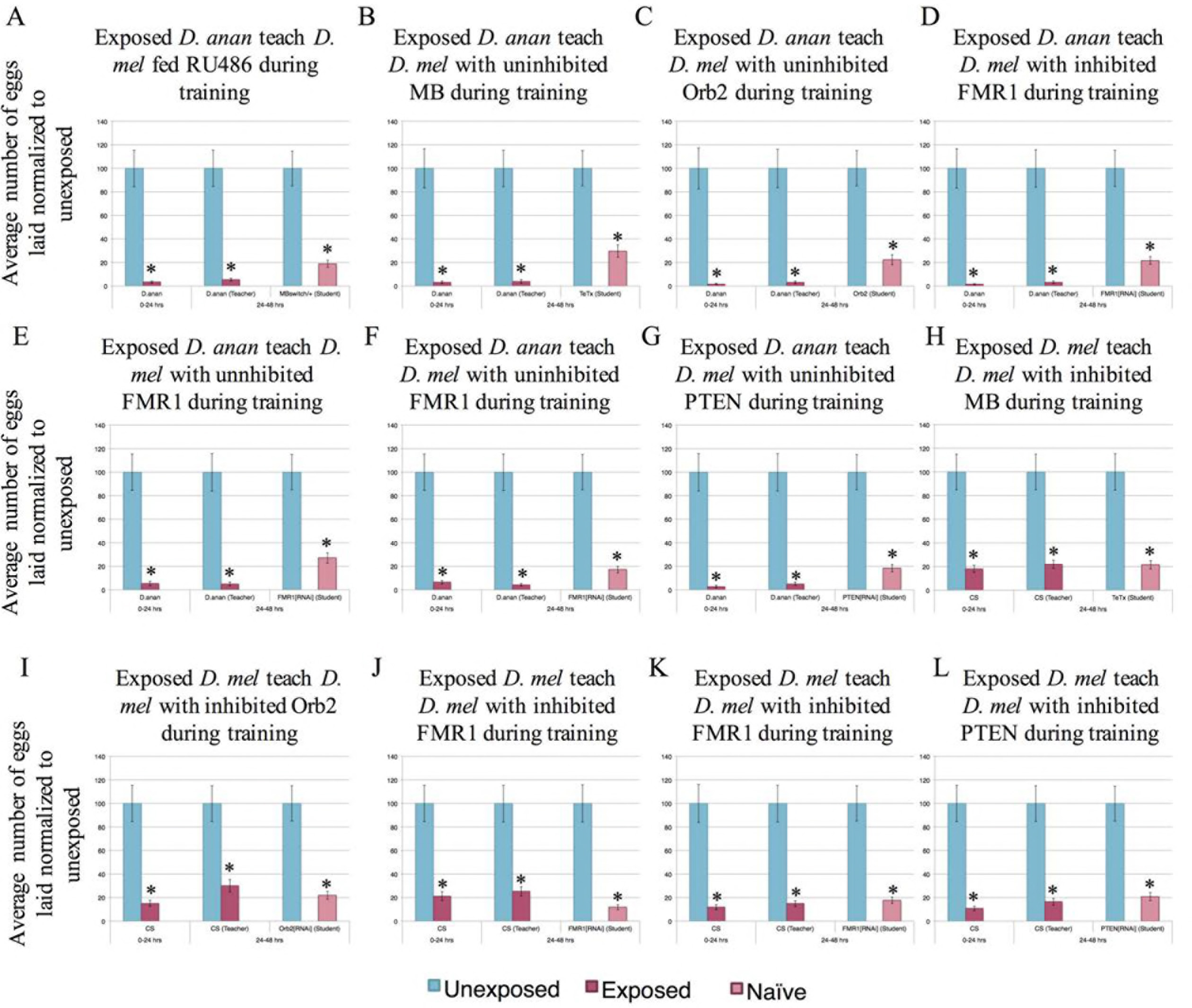
Further evidence showing a critical role for the mushroom body and memory proteins for dialect learning. Percentage of eggs laid by exposed flies normalized to eggs laid by unexposed flies is shown. Communication between trained *D. melanogaster*, MBswitch/+ (outcrossed to *Canton S*) students and *D. ananassae* teachers fed RU486 during the training period shows communication between the two species, demonstrating that RU486 feeding does not perturb dialect learning (A). Communication between trained students *D. melanogaster* and *D. ananassae*, with training by flies not expressing tetanus toxin (UAS-TeTx) in the mushroom body (MB) (i.e. methanol fed), shows communication between the species (B). Communication between *D. ananassae* and students trained with *D. ananassae* with no RNAi-mediated Orb2 knockdown in the MB (i.e. methanol fed) shows communication between the species (C). Communication between *D. ananassae* and students trained with *D. ananassae* with RNAi-mediated FMR1 knockdown (strain #34944) in the MB (i.e. RU486 fed) shows that FMR1 is not required in the MB during the training period (D). Communication between *D. ananassae* and students trained with *D. ananassae* with no FMR1 knockdown (strain #24944) in the MB (i.e. methanol fed) shows wild-type behavior (E). Communication between *D. ananassae* and students trained with *D. ananassae* with no FMR1 knockdown (strain #24944) in the MB (i.e. methanol fed) shows wild-type behavior (F). Communication between *D. ananassae* and students trained with *D. ananassae* with no PTEN knockdown in the MB (i.e. methanol fed) shows wild-type behavior (G). Error bars represent standard error (n = 12 biological replicates) (*p < 0.05). Communication between various *D. melanogaster* lines trained by *D. ananassae* show wild-type communication with *D. melanogaster* (*Canton S*). Lines shown are MB switch expressing TeTx (H), Orb2^RNAi^ (I), FMR1^RNAi^ (strain number 27484) (J), FMR1^RNAi^ (strain number 34944) (K), PTEN^RNAi^ (L), and were fed RU486 during cohabitation with *D. ananassae*.

**Supplementary Figure 13.**
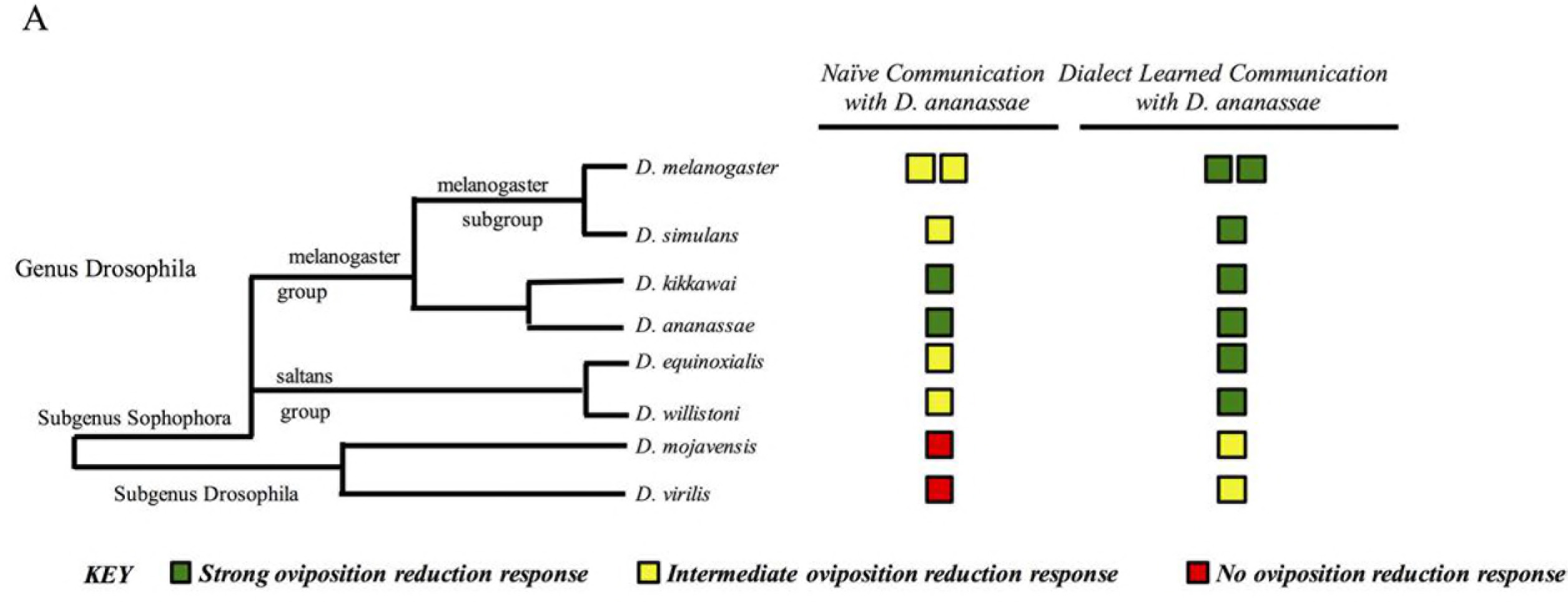
Phylogenetic summary of dialect learning for *D. ananassae*. We utilize species across the genus Drosophila to show communication ability of *D. ananassae* (A). We observe the ability to demonstrate interspecies communication, which varies across the genus, with species closely related to *D. ananassae* able to communicate without barriers. More distantly related species have difficulty communicating, though the barrier can be alleviated with dialect training. Double boxes in a given row and column indicate multiple wild-type strains were tested.

### SUPPLEMENTARY TABLE LEGENDS

**Supplementary Table 1.** Fly lines and species used in this study.

## MATERIALS AND METHODS

### Insect Species/Strains

The *D. melanogaster* strains Canton-S (CS), Oregon-R (OR), white^1118^(*w*^*1118*^), and transgenic flies carrying Histone H2AvD-RFP (His-RFP) were used as wild-type strains. Experiments were primarily performed using CS as wild type flies except where otherwise indicated. *Orco*^*1*^(*OrS3b*^*1*^), UAS-TeTx, UAS-Orb2^RNAi^, UAS-FMR1^RNAi^, UAS-FMR1^RNAi^, UAS-PTEN^RNAi^, UAS-Ir8a^RNAi^, UAS-Ir25a^RNAi^, *ninaB*^*P315*^ were acquired from the Bloomington Drosophila Stock Center (stock numbers 23129, 28838, 27050, 27484, 34944, 25841, 25813, 43985, and 24776 respectively). Drosophila species were acquired from the Drosophila Species Stock Center (DSSC) at the University of California, San Diego. Flies and their respective stock numbers are listed: *D. simulans* (14021-0251.196), *D. mauritiana* (14021-0241.01), *D. sechellia* (14021-0248.25), *D. yakuba* (14021-0261.01), *D. tsacasi* (14028-0701.00), *D. kikkawai* (14028-0561.00), *D. ananassae* (14024-0371.13 and 14024-0371.11), *D. pseudoobscura* (14011-0121.00), *D. neocordata* (14041-0831.00), *D. equinoxialis* (14030-0741.00), *D. willistoni* (14030-0811.00), *D. immigrans* (15111-1731.08), *D. mojavensis* (15081-1352.22), and *D. virilis* (15010-1051.87). All experiments with *D. ananassae* used strain number 14024-0371.13 unless otherwise noted (Table S 1).

The *ewg*^*NS4*^ mutant line was kindly provided by Yashi Ahmed (Geisel School of Medicine at Dartmouth). The mushroom body Gene-Switch line was kindly provided by Greg Roman (Baylor College of Medicine). *IrSa*^*1*^, *Ir25a*^*2*^, *IrSa*>*GAL4*, *Ir25a*>*GAL4* and *IrSa*^*1*^*;Ir25a*^*2*^*;Orco*^*1*^ lines were kindly provided by Greg S. B. Suh (Skirball Institute at NYU). Flies aged 3-6 days post-eclosion on fresh Drosophila media were used in all experiments. Flies were maintained at room temperature with approximately 30% humidity. All species and strains used were maintained in fly bottles (Genesse catalog number 32-130) containing 50 mL of standard Drosophila media. Bottles were supplemented with 3 Kimwipes rolled together and placed into the center of the food. Drosophila media was also scored to promote oviposition. Fly species stocks were kept separate to account for visual cues that could be conferred if the stocks were kept side-by-side.

The Figitid larval endoparasitoid *Leptopilina heterotoma* (strain Lh14) was used in all experiments. *L. heterotoma* strain Lh14 originated from a single female collected in Winters, California in 2002. In order to propagate wasp stocks, we used adult *D. virilis* in batches of 40 females and 15 males per each vial (Genesse catalog number 32-116). Adult flies were allowed to lay eggs in standard Drosophila vials containing 5 mL standard Drosophila media supplemented with live yeast (approximately 25 granules) for 4-6 days before being replaced by adult wasps, using 15 female and 6 male wasps, for infections. These wasps deposit eggs in developing fly larvae, and we gave them access specifically to the L2 stage of *D. virilis* larvae. Wasp containing vials were supplemented with approximately 500 μL of a 50% honey/water solution applied to the inside of the cotton vial plugs. Organic honey was used as a supplement. Wasps aged 3-7 days post eclosion were used for all infections and experiments. Wasps were never reused for experiments.

### Fly Duplexes

Briefly, fly duplexes were constructed (Desco, Norfolk, MA) by using three standard 25mm × 75mm pieces of acrylic that were adhered between two 75mm × 50mm × 3mm pieces of acrylic. Clear acrylic sealant was used to glue these pieces together, making two compartments separated by one 3mm thick acrylic piece. Following sealant curing, each duplex was soaked in water and Sparkleen detergent (Fisherbrand^TM^ catalog number 04-320-4) overnight, then soaked in distilled water overnight and finally air-dried. The interior dimensions of each of the two units measured approximately 23.5mm (wide) × 25mm (deep) × 75mm (tall).

For experiments using Fly Duplexes (teacher-student interaction), bead boxes (6 slot jewelers bead storage box watch part organizer sold by FindingKing) were used to accommodate 12 replicates of each treatment group. Each compartment measures 32 × 114 mm with the tray in total measuring 21 × 12 × 3.5 mm. Each compartment holds 2 duplexes, and the tray in total holds 12 duplexes. Empty duplexes were placed into the bead box compartments. 50 mL standard Drosophila media in a standard Drosophila bottle (Genesse catalog number 32-130) was microwaved for 39 seconds. This heated media was allowed to cool for 2 minutes on ice before being dispensed. Each duplex unit was then filled with 5 mL of the media and further allowed to cool until solidification. The open end of the Fly Duplex was plugged with a cotton plug (Genesse catalog number 51-102B) to prevent insect escape. 10 female flies and 2 male flies were placed into one chamber of the Fly Duplex in the control, while 20 female Lh14 wasps were placed next to the flies in the experimental setting for 24 hours. After the 24-hour exposure, flies and wasps were removed by anesthetizing flies and wasps in the Fly Duplexes. Control flies underwent the same anesthetization. Wasps were removed and replaced with 10 female and two male “student” flies. All flies were placed into new clean duplexes for the second 24-hour period, containing 5 mL Drosophila media in a new bead box. For fly duplexes containing a subset of species, specifically *D. mojavensis*, *D. immigrans*, and *D. virilis*, 10 yeast granules were added to the standard Drosophila media after solidification of the food. This activated yeast was added to promote oviposition. Flies showed minimal oviposition in food lacking yeast. We speculate this was observed due to the fly food being optimized for *D. melanogaster*. Plugs used to keep insects in the duplex were replaced every 24 hours to prevent odorant deposition on plugs that could influence behavior. The oviposition bead box from each treatment was replaced 24 hours after the start of the experiment, and the second bead box was removed 48 hours after the start of the experiment. Fly egg counts from each bead box were made at the 0-24 and 24-48-hour time points.

All experimental treatments were run at 25°C with a 12:12 light:dark cycle at light intensity 16_7_, using twelve replicates at 40% humidity unless otherwise noted. Light intensity was measured using a Sekonic L-308DC light meter. The light meter measures incident light and was set at shutter speed 120, sensitivity at iso8000, with a 1/10 step measurement value (f-stop). Fly duplexes and bead boxes soaked with distilled water mixed with Sparkleen after every use for 4 hours at minimum and subsequently rinsed with distilled water and air-dried. All egg plates were coded and scoring was blind as the individual counting eggs was not aware of treatments or genotypes.

### Dialect Exposure

Species were cohabitated in standard Drosophila bottles (Genesee catalog number 32-130) containing 50 mL standard Drosophila media. Three Kimwipes were rolled together and placed into the center of the food. Batches of 3 bottles were made per treatment. Two species were incubated in each bottle with 100 female and 20 males of each species per bottle. Every two days, flies were placed into new bottles prepared in the identical manner. Flies were cohabitation for approximately 168 hours (7 days), unless otherwise noted. Following cohabitation, flies were anesthetized and the two species were separated. The flies were then used as students to wasp or mock exposure teachers of the opposite species. For example, we cohabitated *D. melanogaster* and *D. ananassae* for one week. Following the weeklong cohabitation, we separated the dialect-trained flies. Trained *D. melanogaster* were placed in duplexes next to *D. ananassae* either mock or wasp exposed. Trained *D. ananassae* were placed in duplexes next to *D. melanogaster* either mock treated or wasp exposed.

For experiments utilizing more than two species for dialect learning, species were cohabitated in standard Drosophila bottles (Genesee catalog number 32-130) containing 50 mL standard Drosophila media. Three Kimwipes were rolled together and placed into the center of the food. Batches of 3 bottles were made per treatment. The three species were incubated in each bottle with 100 female and 20 males of each species per bottle. Every two days, flies were placed into new bottles prepared in the identical manner. The three-fly species were cohabitation for approximately 168 hours (7 days), unless otherwise noted. Following cohabitation, flies were anesthetized and one of the three species was tested by pairing them with teachers of the other two species. For example we cohabitated *D. melanogaster*, *D. ananassae*, and *D. willistoni* for one week. Following the weeklong cohabitation, we separated the dialect-trained flies. Trained *D. melanogaster* were placed in duplexes next to either *D. ananassae* or *D. willistoni*, mock or wasp exposed.

For cohabitation experiments where two species were allowed visual only cues, the Fly Duplex was utilized. The two species were co-incubated side-by-side with 100 female and 20 males of each species per unit using the two chambers of the fly duplex such that the flies could only see each other. The fly duplex was placed into bead boxes, with each unit of the duplex containing 5 mL of standard Drosophila media. Every two days, flies were placed into new fly duplexes with fresh 5 mL standard Drosophila media. Following the weeklong co-incubation, flies were anesthetized and the two species were separated. The flies were then used as students to wasp or mock exposure teachers of the opposite species.

For cohabitation experiments where the two species did not have visual cues, the two species were incubated in bottles with 100 female and 20 males of each species per bottle in complete darkness. The only difference between this method and other training sessions was the lack of light—meaning flies were subject to 25°C with 40% humidity. Every two days, flies were placed into new bottles prepared in the identical manner. Flies were exposed to light for less than 30 seconds, during which they were placed into a new bottle, and immediately returned to the dark. Following the weeklong dark-cohabitation, flies were anesthetized and the two species were separated. The flies were then used as students to wasp or mock exposure teachers of the opposite species.

For cohabitation experiments under monochromatic light settings, batches of 3 bottles with 100 female and 20 males of each species were placed into 27.9cm × 16.8cm × 13.7cm plastic boxes (Sterilite 1962 Medium Clip Box with Blue Aquarium Latches sold by Flikis). These boxes were externally wrapped with colored cellophane wrap, allowing only a certain wavelength of light to be transmitted into the boxes. Red and blue cellophane wraps were purchased from Amscam (Amscan Party Supplies for Any Occasion Functional Cellophane Wrap, 16' × 30", Rose Red and Spanish Blue). Cellophane wrapped boxes with bottles containing flies were subject to 25°C with 40% humidity under the same light intensity as previous experiments. Light intensity within the red wrapped box was 11_2_ and within the blue wrapped box was 11_5_ measured using the Sekonic L-308DC light meter. Every two days, flies were placed into new bottles prepared in the manner described previously. Flies were exposed to broad-spectrum light for less than 30 seconds, during which they were placed into a new bottle, and immediately returned to monochromatic light. Following the weeklong monochromatic-light-cohabitation, flies were anesthetized and the two species were separated. The flies were then used as students to wasp or mock exposure teachers of the opposite species.

For the one-day cohabitation experiments, batches of 3 bottles with 100 female and 20 males of each species were placed at 25°C with 40% humidity for 24 hours. Following the 24-hour cohabitation, flies were anesthetized and the two species were separated. The flies were then used as students to wasp or mock exposure teachers of the opposite species.

### RU486 feeding

RU486 (Mifepristone) was used from Sigma (Lot number SLBG0210V) as the ligand for Gene-Switch experiments. Dialect training bottles were prepared by directly pipetting an RU486 solution onto the 3 Kimwipes in the bottle. The solution was prepared by dissolving 3.575 mg of RU486 in 800μL methanol (Fisher Scientific Lot number 141313). This solution was added to 15.2 mL of distilled water. The total solution (16 mL) was thoroughly mixed and 4000 μL was pipetted onto the Kimwipe in each bottle. For bottles containing no RU486 (methanol only) 800μL methanol was mixed with 15.2 mL of distilled water. The total solution (16 mL) was thoroughly mixed and 4000 μL were pipetted onto the Kimwipe in each bottle. Flies were shifted to new bottles prepared in the exact same manner every two days. Flies were cohabitated for approximately 7 days. Following cohabitation, flies were anesthetized and the two species were separated. The flies were then used as students to wasp or mock exposure teachers of the opposite species.

### Immunofluorescence

Ovaries were collected from flies that were placed in vials along with female wasps for experimental or no wasps for control settings. Flies were placed in batches into standard vials (Genesee catalog number 32-116) of 20 females, 2 males along with 20 female wasps for exposed vials, or simple placing 20 female and 2 male flies in vials for the unexposed treatments. Three vials were prepared to produce three replicates to account for batch effects. We observed no batch effects so each of the 12 ovaries imaged from each treatment were then counted as a replicate, thus providing an n of 36. Ovaries that were prepared for immunofluorescence were fixed in 4% methanol-free formaldehyde in PBS with 0.001% Triton-X for approximately five minutes. The samples were then washed in PBS with 0.1% Triton-X, and blocked with 2% normal goat serum (NGS) for two hours. The primary antibody, cleaved Drosophila Dcp-1 (Asp216) (Cell Signaling number 9578) at a concentration of 1:100, was used to incubate the ovaries overnight at 4° C in 2% normal goat serum (NGS). The secondary antibody used was Fluorescein isothiocyanate (FITC) conjugated (Jackson Immunoresearch), and used at a concentration of 1:200 for a two-hour incubation at room temperature. This was followed by a 10-minute nuclear stain with 4', 6-diamidino-2-phenylindole (DAPI). For confocal imaging of *D. melanogaster* ovaries, wheat germ agglutinin (WGA) was also used as a membrane marker (Fig. 1 F,J, Fig. S 2).

### Imaging

A Nikon A1R SI Confocal microscope was used for imaging activated Dcp-1 caspase staining in *D. melanogaster* (Fig. 1 D-K, Fig. S 2). Image averaging of 4x during image capture was used for all images. A Nikon E800 Epifluorescence microscope with Olympus DP software was used to image Dcp-1 caspase staining on all other Drosophila species tested. This microscope was also used to quantify egg chambers with Dcp-1 signal and total number of egg chambers in all species tested.

### Statistical analysis

Statistical tests were performed in Microsoft Excel. Welch’s two-tailed t-tests were performed for data. P-values reported were calculated for comparisons between paired treatment-group and unexposed.

## REFERENCES

(1) Gould JL. Honey bee communication. Nature 1974.

(2) Wenner AM. Sound production during the waggle dance of the honey bee. Anim Behav 1962; 10(1):79–95.

(3) Winston ML. The biology of the honey bee. : harvard university press; 1991.

(4) Goodale E, Beauchamp G, Magrath RD, Nieh JC, Ruxton GD. Interspecific information transfer influences animal community structure. Trends in ecology & evolution 2010;25(6):354–361.

(5) Westrip JR, Bell MB. Breaking down the species boundaries: selective pressures behind interspecific communication in vertebrates. Ethology 2015;121(8):725–732.

(6) Elgar MA, Nash DR, Pierce NE. Eavesdropping on cooperative communication within an ant-butterfly mutualism. The Science of Nature 2016;103(9–10):84.

(7) Virant-Doberlet M, Mazzoni V, De Groot M, Polajnar J, Lucchi A, Symondson WO, et al. Vibrational communication networks: eavesdropping and biotic noise. Studying vibrational communication: Springer; 2014.p.93–123.

(8) Lima SL. Predators and the breeding bird: behavioral and reproductive flexibility under the risk of predation. Biological reviews 2009;84(3):485–513.

(9) Harbison H, Nelson DA, Hahn TP. Long-term persistence of song dialects in the mountain white-crowned sparrow. Condor 1999:133–148.

(10) Koetz AH, Westcott DA, Congdon BC. Spatial pattern of song element sharing and its implications for song learning in the chowchilla, Orthonyx spaldingii. Anim Behav 2007;74(4):1019–1028.

(11) Marler P, Tamura M. Culturally Transmitted Patterns of Vocal Behavior in Sparrows. Science 1964 Dec 11;146(3650):1483–1486.

(12) Soha JA, Nelson DA, Parker PG. Genetic analysis of song dialect populations in Puget Sound white-crowned sparrows. Behav Ecol 2004;15(4):636–646.

(13) Baptista LF, Morton ML. Song learning in montane white-crowned sparrows: from whom and when. Anim Behav 1988;36(6): 1753–1764.

(14) Alem S, Perry CJ, Zhu X, Loukola OJ, Ingraham T, Søvik E, et al. Associative mechanisms allow for social learning and cultural transmission of string pulling in an insect. PLoS Biol 2016;14(10):e1002564.

(15) Loukola OJ, Perry CJ, Coscos L, Chittka L. Bumblebees show cognitive flexibility by improving on an observed complex behavior. Science 2017 Feb 24;355(6327):833–836.

(16) Battesti M, Moreno C, Joly D, Mery F. Spread of social information and dynamics of social transmission within Drosophila groups. Current Biology 2012;22(4):309–313.

(17) Kacsoh BZ, Bozler J, Ramaswami M, Bosco G. Social communication of predator-induced changes in Drosophila behavior and germ line physiology. Elife 2015 May 13;4:10.7554/eLife.07423.

(18) Beruter J, Beauchamp GK, Muetterties EL. Complexity of chemical communication in mammals: Urinary components mediating sex discrimination by male guinea pigs. Biochem Biophys Res Commun 1973; 53(1):264–271.

(19) Apfelbach R, Parsons MH, Soini HA, Novotny MV. Are single odorous components of a predator sufficient to elicit defensive behaviors in prey species? Front Neurosci 2015 Jul 29;9:263.

(20) Voznessenskaya VV. Influence of cat odor on reproductive behavior and physiology in the house mouse (Mus musculus). Neurobiology of Chemical Communication (Frontiers in Neuroscience Book Series), C. Musignat-Caretta (Ed), CRC Press 2014:389–405.

(21) Schuchmann M, Siemers BM. Behavioral evidence for community-wide species discrimination from echolocation calls in bats. Am Nat 2010;176(1):72–82.

(22) Herzingl D, Johnsonz C. Interspecific interactions between Atlantic spotted dolphins (Stenella fr0ntalis)'and bottlenose dolphins (T ursiops truncatus) in the. Aquat Mamm 1997;23:85–99.

(23) Meinke DW, Cherry JM, Dean C, Rounsley SD, Koornneef M. Arabidopsis thaliana: a model plant for genome analysis. Science 1998;282(5389):662–682.

(24) Heil M, Karban R. Explaining evolution of plant communication by airborne signals. Trends in ecology & evolution 2010;25(3):137–144.

(25) Bruin J, Dicke M, Sabelis M. Plants are better protected against spider-mites after exposure to volatiles from infested conspecifics. Experientia 1992;48(5):525–529.

(26) Karban R, Shiojiri K, Ishizaki S. An air transfer experiment confirms the role of volatile cues in communication between plants. Am Nat 2010; 176(3):381–384.

(27) Karban R, Baldwin IT. Induced responses to herbivory. : University of Chicago Press; 2007.

(28) Karban R, Baldwin I, Baxter K, Laue G, Felton G. Communication between plants: induced resistance in wild tobacco plants following clipping of neighboring sagebrush. Oecologia 2000;125(1):66–71.

(29) Kost C, Heil M. Herbivore-induced plant volatiles induce an indirect defence in neighbouring plants. J Ecol 2006;94(3):619–628.

(30) Rhoades DF. Responses of alder and willow to attack by tent caterpillars and webworms: evidence for pheromonal sensitivity of willows.; 1983.

(31) Shulaev V, Silverman P, Raskin I. Airborne signalling by methyl salicylate in plant pathogen resistance.[Erratum: Apr 17, 1997, v. 386 (6626), p. 738.]. Nature 1997.

(32) Karban R, Shiojiri K, Huntzinger M, McCall AC. Damage-induced resistance in sagebrush: volatiles are key to intra-and interplant communication. Ecology 2006;87(4):922–930.

(33) Farmer EE, Ryan CA. Interplant communication: airborne methyl jasmonate induces synthesis of proteinase inhibitors in plant leaves. Proceedings of the National Academy of Sciences 1990;87(19):7713–7716.

(34) Glinwood R, Ninkovic V, Pettersson J, Ahmed E. Barley exposed to aerial allelopathy from thistles (Cirsium spp.) becomes less acceptable to aphids. Ecol Entomol 2004;29(2):188–195.

(35) Fowler SV, Lawton JH. Rapidly induced defenses and talking trees: the devil's advocate position. Am Nat 1985:181–195.

(36) Bier E. Drosophila, the golden bug, emerges as a tool for human genetics. Nature Reviews Genetics 2005;6(1):9–23.

(37) Helfand SL, Rogina B. Genetics of aging in the fruit fly, Drosophila melanogaster. Annu Rev Genet 2003;37(1):329–348.

(38) Markow TA. The secret lives of Drosophila flies. Elife 2015 Jun 4;4:10.7554/eLife.06793.

(39) Markow T, Beall S, Castrezana S. The wild side of life: reproductive biology of Drosophila in nature. Fly 2012;6:98–101.

(40) Markow TA. “Cost” of virginity in wild Drosophila melanogaster females. Ecology and evolution 2011;1(4):596–600.

(41) Markow TA. Forced Matings in Natural Populations of Drosophila. Am Nat 2000 Jul;156(1):100–103.

(42) Schlenke TA, Morales J, Govind S, Clark AG. Contrasting infection strategies in generalist and specialist wasp parasitoids of Drosophila melanogaster. PLoS Pathog 2007;3(10):e158.

(43) Kacsoh BZ, Bozler J, Hodge S, Ramaswami M, Bosco G. A novel paradigm for nonassociative long-term memory in Drosophila: predator-induced changes in oviposition behavior. Genetics 2015 Apr;199(4): 1143–1157.

(44) Kacsoh BZ, Lynch ZR, Mortimer NT, Schlenke TA. Fruit flies medicate offspring after seeing parasites. Science 2013 Feb 22;339(6122):947–950.

(45) Lefevre T, de Roode JC, Kacsoh BZ, Schlenke TA. Defence strategies against a parasitoid wasp in Drosophila: fight or flight? Biol Lett 2012 Apr 23;8(2):230–233.

(46) Lynch ZR, Schlenke TA, Roode J. Evolution of behavioural and cellular defences against parasitoid wasps in the Drosophila melanogaster subgroup. J Evol Biol 2016.

(47) Van Der Linde K, Houle D, Spicer GS, Steppan SJ. A supermatrix-based molecular phylogeny of the family Drosophilidae. Genetics research 2010;92(1):25.

(48) DeSimone S, Coelho C, Roy S, VijayRaghavan K, White K. ERECT WING, the Drosophila member of a family of DNA binding proteins is required in imaginal myoblasts for flight muscle development. Development 1996 Jan;122(1):31–39.

(49) Larsson MC, Domingos AI, Jones WD, Chiappe ME, Amrein H, Vosshall LB. Or83b encodes a broadly expressed odorant receptor essential for Drosophila olfaction. Neuron 2004;43(5):703–714.

(50) Benton R, Vannice KS, Gomez-Diaz C, Vosshall LB. Variant ionotropic glutamate receptors as chemosensory receptors in Drosophila. Cell 2009;136(1):149–162.

(51) Mao Z, Roman G, Zong L, Davis RL. Pharmacogenetic rescue in time and space of the rutabaga memory impairment by using Gene-Switch. Proc Natl Acad Sci U S A 2004 Jan 6;101(1): 198–203.

(52) Keleman K, Krüttner S, Alenius M, Dickson BJ. Function of the Drosophila CPEB protein Orb2 in long-term courtship memory. Nat Neurosci 2007;10(12):1587–1593.

(53) Chiang HC, Wang L, Xie Z, Yau A, Zhong Y. PI3 kinase signaling is involved in Abeta-induced memory loss in Drosophila. Proc Natl Acad Sci U S A 2010 Apr 13;107(15):7060–7065.

(54) Silverman JL, Yang M, Lord C, Crawley JN. Behavioural phenotyping assays for mouse models of autism. Nature Reviews Neuroscience 2010;11(7):490–502.

(55) Shevtsova E, Hansson C, Janzen DH, Kjaerandsen J. Stable structural color patterns displayed on transparent insect wings. Proc Natl Acad Sci U S A 2011 Jan 11;108(2):668–673.

(56) Katayama N, Abbott JK, Kjaerandsen J, Takahashi Y, Svensson EI. Sexual selection on wing interference patterns in Drosophila melanogaster. Proc Natl Acad Sci U S A 2014 Oct 21;111(42):15144–15148.

